# Sox2 levels configure the WNT response of epiblast progenitors responsible for vertebrate body formation

**DOI:** 10.1101/2020.12.29.424684

**Authors:** Robert Blassberg, Harshil Patel, Thomas Watson, Mina Gouti, Vicki Metzis, M Joaquina Delás, James Briscoe

## Abstract

WNT signalling has multiple roles. It maintains pluripotency of embryonic stem cells, assigns posterior identity in the epiblast and induces mesodermal tissue. We provide evidence that these distinct functions are conducted by the transcription factor SOX2, which adopts different modes of chromatin interaction and regulatory element selection depending on its level of expression. At high levels, SOX2 acts as a pioneer factor, displacing nucleosomes from regulatory elements with high affinity SOX2 binding sites and recruiting the WNT effector, TCF/β-catenin, to maintain pluripotent gene expression. Reducing SOX2 levels destabilises pluripotency and reconfigures SOX2/TCF/β-catenin occupancy to caudal epiblast expressed genes. These contain low-affinity SOX2 sites and are co-occupied by T/Bra and CDX. The loss of SOX2 allows WNT induced mesodermal differentiation. These findings define a role for Sox2 levels in dictating the chromatin occupancy of TCF/β-catenin and reveal how context specific responses to a signal are configured by the level of a transcription factor.

## Introduction

The development and maintenance of the diversity of cell types that comprise a multicellular organism is achieved through the spatial and temporal regulation of gene expression, controlled and organised by extrinsic signals. But there are relatively few signals compared to the number and variety of cell types and these signals are reused multiple times over the course of ontogeny. This raises the question of how individual signals produce a diversity of outputs. While it is clear that alterations in the molecular composition of cells during development plays a key role in determining how cells respond to extrinsic signals, the molecular mechanisms that ensure context-dependent responses remain incompletely understood.

An example of this is WNT signalling which, through its transcriptional effector TCF/β-catenin, has multiple functions in development (Steinhart and Angers, 2018) (Cadigan and Waterman, 2012). In mouse embryonic stem (mES) cells, WNT/β-catenin signalling promotes pluripotency (Madeja et al., 2015) (Ying et al., 2008). As development proceeds, WNT/β-catenin signalling assigns posterior positional identity to regionalize the embryo by upregulating the expression of CDX transcription factors (TFs) in the forming caudal epiblast (Deschamps and Nes, 2005). A subset of these cells, located within the caudal lateral epiblast (CLE), are neuromesodermal progenitors (NMPs) that generate the neural and mesodermal tissue responsible for the elongation of the axis (Henrique et al., 2015). In NMPs, WNT/β-catenin signalling promotes differentiation to mesodermal tissue at the expense of spinal cord neural differentiation (Martin and Kimelman, 2012) (Tsakiridis et al., 2014) (Gouti et al., 2014)(Garriock et al., 2015)(Gouti et al., 2017)(Koch et al., 2017)(Veenvliet et al., 2020).

Alongside WNT signalling, the TF SOX2 also plays a central role. SOX2 is expressed at high levels in mES cells and epiblast cells where it is necessary to maintain the undifferentiated pluripotent state (Avilion et al., 2003) (Masui et al., 2007). Then, as the embryo regionalizes, SOX2 expression drops in the CLE and remains expressed at low levels in NMPs together with genes conferring primitive streak identity such as T/Bra (Wood and Episkopou, 1999) (Wymeersch et al., 2016) (Mulas et al., 2018). Upregulation of SOX2 is associated with the allocation of NMPs to spinal cord neural progenitors (Gouti et al., 2014) (Kinney et al., 2020). By contrast, SOX2 expression is lost upon commitment of NMPs to mesodermal lineages (Wymeersch et al., 2016). This involves repression by the TF TBX6. Genetic ablation of *Tbx6* results in the upregulation of SOX2 expression in paraxial mesodermal progenitors and the differentiation of these cells to ectopic neural structures (Takemoto et al., 2011). In vitro assays indicate that TBX6 null progenitors cultured under caudal epiblast inducing conditions harbour low levels of SOX2 and are maintained in an undifferentiated state that acquires an increasingly posterior identity in the absence of differentiation to paraxial mesoderm (Gouti et al., 2017). This implicates low levels of SOX2 in coordinating the maintenance of neuromesodermal potential and patterning of the embryonic axis.

The correlation between SOX2 levels and the changes in the function of WNT signalling raises the possibility that SOX2 influences the response of cells to WNT signalling. In mES cells, SOX2 acts with β-catenin and the TFs TCF7L1, OCT4 and NANOG to promote the expression of WNT target genes that maintain the undifferentiated pluripotency state (Boyer et al., 2005) (Cole et al., 2008) (Yi et al., 2011). By contrast, genomic analysis of in vitro differentiated NMPs revealed a large number of cis-regulatory elements (CREs) associated with mesodermal genes that are co-occupied by SOX2, T/BRA and TBX6 and B-catenin (Koch et al., 2017). This led to the suggestion that SOX2 contributes to maintaining the undifferentiated state of CLE progenitors by directly counteracting WNT signalling activity and inhibiting mesoderm gene expression. Consistent with this, SOX3, the closely related paralogue of SOX2, physically interacts with β-catenin and inhibits its activity (Zorn et al., 1999). Nevertheless, direct evidence of whether and how SOX2 is responsible for the different developmental responses to WNT signalling has been lacking.

To test the causal role of SOX2 in the response of cells to WNT signalling, we decoupled SOX2 expression from its developmental regulation. This revealed that SOX2 directly controls the WNT response by adopting different modes of chromatin interaction and occupying distinct genomic locations depending on its level of expression. We provide evidence that the arrangement of SOX2 binding regulates the genomic occupancy of TCF/β-catenin and determines the expression of cell-type specific WNT target genes. High levels of SOX2 maintain chromatin accessibility and TCF/β-catenin occupancy at CREs associated with pluripotency genes. Downregulation of SOX2 leads to loss of accessibility at these CREs and a shift in SOX2/TCF/β-catenin complexes to CREs associated with caudal epiblast and paraxial mesoderm genes. This alters the transcriptional response to WNT signalling from a pluripotency to a CLE programme. Occupancy at caudal epiblast specific genes is not determined by changes in chromatin accessibility. Instead it correlates with expression of CDX and T/BRA TFs. SOX2 directly promotes expression of CDX2 in response to WNT signalling in caudal epiblast progenitors, and this interaction is modulated by a feedback loop between SOX2 and Nodal signalling. We propose that the low level of SOX2 expression in caudal epiblast progenitors underlies their distinct transcriptional identity by both promoting expression of caudal-epiblast specific WNT-responsive targets and disabling the expression of pluripotency genes that inhibit differentiation. Together these results provide insight into the mechanisms that determine the context specific response of cells to an extrinsic signal that generates the diversity of outcomes necessary for tissue development.

## Results

### Altering Sox2 levels changes the response of pluripotent cells to WNT signaling

To investigate how SOX2 functions in the CLE, we took advantage of an in vitro model of the caudal epiblast. Mouse ESCs differentiated for 48h in the presence of FGF and LGK974, an inhibitor of WNT secretion (henceforth ‘FL-media’), acquire an epiblast-like cell identity (EpiLC) (Fig 1A), recapitulating gene expression changes characteristic of the post-implantation epiblast (Fig S1A). Similar to their *in vivo* counterparts, EpiLCs acquired caudal epiblast-like cell (CEpiLC) identity in response to WNT signalling, activated by 24h exposure to the GSK3 inhibitor CHIR99021 (henceforth ‘FLC-media’) (Fig 1A). This resulted in reduced expression of SOX2, upregulation of the primitive streak marker T/BRA (Fig S1B) and expression of *Cdx* and posterior Hox genes (Fig S1C,D), as previously shown (Gouti et al., 2014).

**Figure 1.**
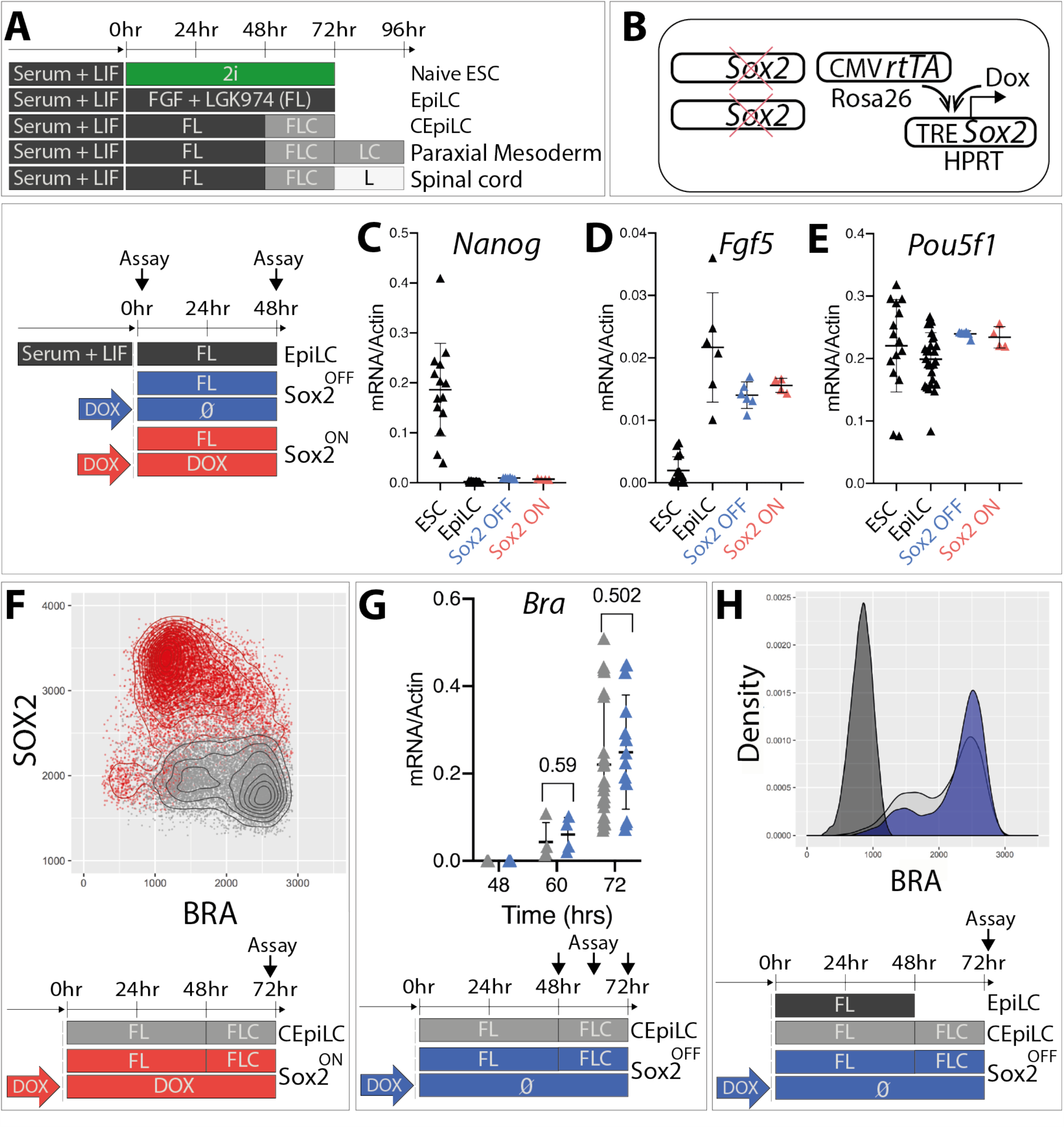
High SOX2 levels inhibit WNT-induced differentiation of epiblast-like cells. (A) Schematic of mESC differentiation. LIF-Leukemia inhibitory factor; F-FGF; L-LGK974; C-CHIR99021; 2i-CHIR99021 + PD173074. (B) To generate SOX2-TetON cells, a Dox-inducible SOX2 transgene was incorporated into the HPRT locus of a mouse embryonic stem-cell line constitutively expressing the rtTA gene from the Rosa26 locus prior to ablation of endogenous SOX2 expression by gene-editing. SOX2-ON and SOX2-OFF cultured in FL-medium downregulate Nanog (C), upregulate FGF5 (D), and maintain Pou5f1 expression (E). (F) SOX2 levels are negatively correlated with BRA expression in SOX2-ON cultured in FLC medium. (G) The kinetics of T/Bra mRNA induction, and (H) the distribution of T/BRA levels is unaltered in SOX2 OFF cultured in FLC medium.

To test the role of SOX2 levels during the specification of CEpiLC identity, we ablated endogenous *Sox2* and introduced a *Sox2* transgene under the control of doxycycline (Fig 1B). We refer to this line as SOX2^TetON^. Addition of doxycycline (Dox) to SOX2^TetON^ generated levels of SOX2 expression similar to those found in wild-type ESCs (Fig S1E) and these cells (henceforth referred to as SOX2-ON) could be propagated as dome-shaped colonies in the presence of Dox in naïve pluripotent “2i” media (Ying et al., 2008) (Fig S1F). Removal of Dox from SOX2^TetON^ (henceforth referred to as SOX2-OFF) resulted in the progressive downregulation of SOX2 (Fig S1G), flattening of colonies (Fig S1H), and loss of expression of pluripotency markers OCT4 and NANOG (Fig S1I). Strikingly, the downregulation of SOX2 led to the induction of T/BRA (Fig S1J).

Differentiation of ESCs to CEpiLC identity involves exiting the pluripotent state and transitioning to EpiLC identity prior to activation of WNT signalling (Gouti et al., 2014) (Fig 1A). Both SOX2-OFF and SOX2-ON differentiated in FL-medium maintained the characteristic pattern of gene expression changes accompanying the transition to EpiLC identity, downregulating *Nanog* and upregulating *Fgf5* while continuing to express *Pou5f1* (Fig 1C-E). Upon transfer to FLC-medium, limited T/BRA induction was observed in SOX2-ON cells and its expression was negatively correlated with SOX2 levels (Fig 1F), consistent with SOX2 acting as a repressor of primitive streak identity (Gouti et al., 2017) (Koch et al., 2017)(Wang et al., 2012)(Thomson et al., 2011). Unexpectedly, in Sox2-OFF cells cultured in FLC-media, in which SOX2 levels drop over a period of 36hrs (Fig S1G), *T/Bra* was induced with the same dynamics (Fig 1G) and resulted in a similar proportion of T/BRA+ cells as WT CEpiLC (Fig 1H).

Analysis of the transcriptome of wildtype (WT) and SOX2-OFF cells differentiated for 24hrs in FLC-media revealed that although genes characteristic of the primitive streak, including T/BRA, were induced in both, a set of genes associated with the caudal epiblast were not upregulated in SOX2-OFF cells (Fig 2A). These included the caudal epiblast determinants *Cdx2* and *Cdx4* and posterior Hox genes. Anterior Hox and paraxial mesoderm marker expression was also reduced (Fig 2A,B, S2A), whereas genes characteristic of earlier more anterior mesoderm and endoderm, including genes involved in endoderm formation and heart development, were increased specifically in SOX2-OFF cells (Fig 2A,C, S2B). Thus, loss of SOX2 disrupts the induction of the caudal epiblast gene expression programme in response to WNT signalling and instead leads to the differentiation of an earlier, more anterior primitive streak identity.

**Figure 2.**
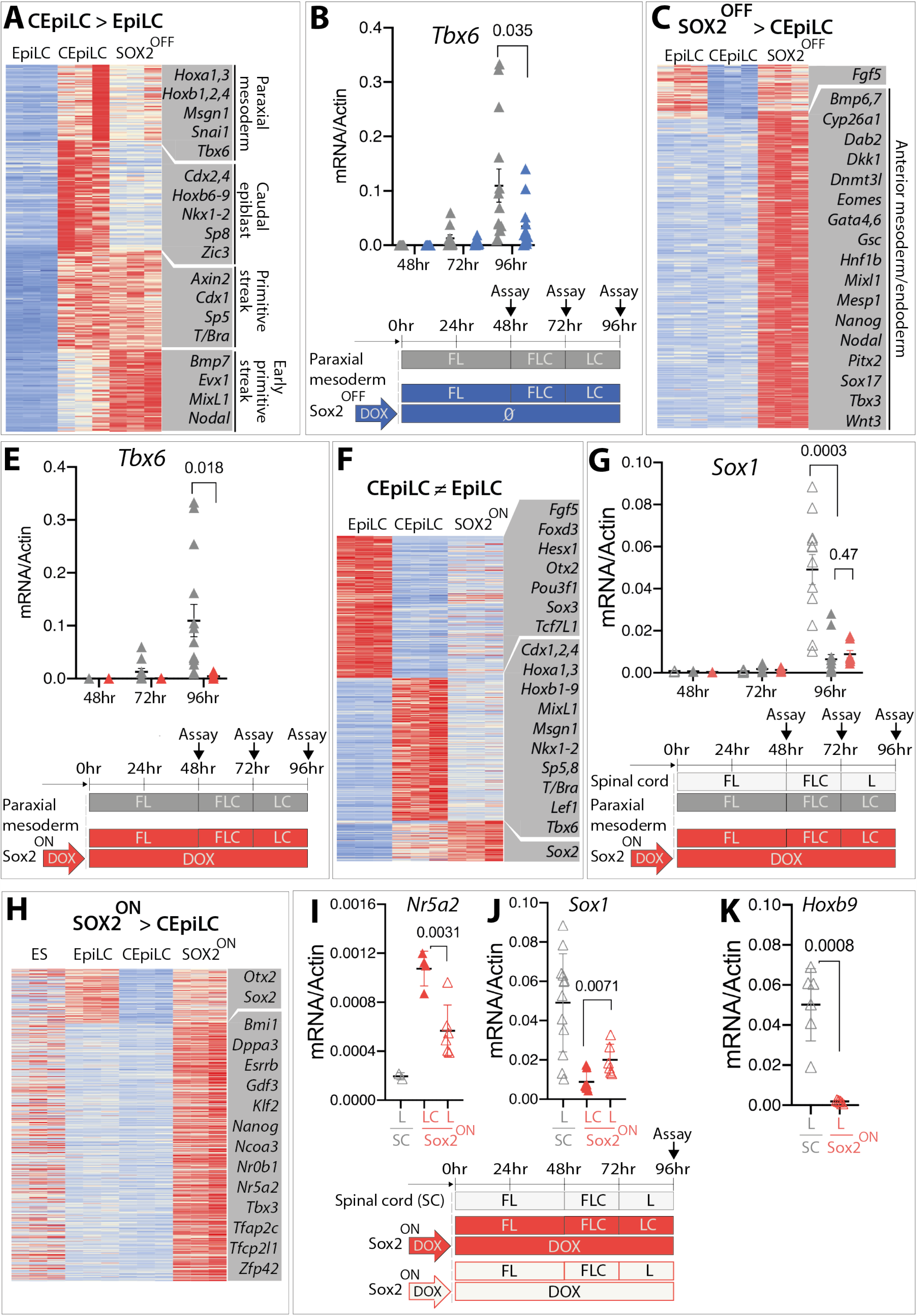
SOX2 dynamics configure the WNT response of epiblast-like cells. (A) Sox2-OFF cultured in FLC medium exhibit altered expression of CEpiLC specific genes. Genes shown are differentially expressed (mod[log2FC] >1, FDR < 0.05) in a comparison between EpiLC and CEpiLC. k-means clustering (k=3). Illustrative genes for each cluster are highlighted. (B) Paraxial mesoderm marker (*Tbx6*) expression is reduced in SOX2-OFF compared to CEpiLCs. (C) Genes upregulated in SOX2-OFF compared to CEpiLCs include early/anterior streak and mesendoderm markers. Genes shown are differentially expressed (log2FC >1, FDR < 0.05) in a comparison between SOX2 OFF and CEpiLC. k-means clustering (k=2). Illustrative genes for each cluster are highlighted. (E) Paraxial mesoderm differentiation is repressed in SOX2-ON cultured in FLC medium. (F) SOX2-ON cultured in FLC medium express low levels of EpiLC and CEpiLC-specific genes. Genes shown are differentially expressed (mod[log2FC] >1, FDR < 0.05) in a comparison between EpiLCs and CEpiLC. k-means clustering (k=3). Illustrative genes for each cluster are highlighted. (G) *Sox1* is not induced in SOX2-ON cultured in FLC medium. (H) SOX2-ON express elevated levels of pluripotency associated genes compared to CEpiLCs and EpiLCs. Genes shown are differentially expressed (log2FC >1, FDR < 0.05) in a comparison between SOX2-ON and CEpiLCs. Illustrative genes for each cluster are highlighted. (I) Expression of the pluripotency factor *Nr5a2* is reduced, and (J) the neural marker S*ox1* increases when SOX2-ON are transferred from FLC to FL medium. (K) The posterior marker *Hoxb9* is not expressed when SOX2-ON are transferred from FLC to FL medium.

We next determined the identity of high SOX2 expressing SOX2-ON cells cultured in CEpiLC differentiation conditions. As predicted from the reduction of T/Bra expression (Fig 1F), paraxial mesoderm differentiation was inhibited (Fig 2E). Moreover, transcriptome analysis revealed that the majority of WNT-induced genes associated with CEpiLC identity were repressed in SOX2-ON cells (Fig 2-F). Contrary to expectations (Koch et al., 2017)(Gouti et al., 2017), SOX2-ON cells cultured in CEpiLC conditions did not differentiate to neural identity (Fig 2G), and instead re-expressed genes associated with naïve pluripotency including *Dppa3, Nanog, Tfap2c*, and *Nr5a2*, (Fig 2H, S2C). Thus, sustaining high levels of SOX2 in the presence of WNT signalling appeared to revert cells to a pluripotent-like state.

We reasoned that removal of the WNT-agonist from SOX2-ON cells should destabilise the pluripotent state and permit their differentiation to neural identity. Consistent with this, a decline in pluripotency marker expression was accompanied by the onset of neural differentiation following WNT-agonist withdrawal from SOX2-ON cells (Fig 2I,J). Importantly, posterior Hox genes, typical of spinal cord neural progenitors, were not induced (Fig 2K), consistent with the idea that downregulation of SOX2 is necessary to disengage the WNT response typical of pluripotent cells and allow the assignment of posterior positional identity prior to trunk lineage differentiation. Taken together these data indicate that a reduction of SOX2 levels is necessary to prevent cells adopting a pluripotent WNT-response but premature elimination of SOX2 abrogates the ability of WNT signalling to promote a caudal identity. This raises the question of how SOX2 alters the response of epiblast progenitors to WNT signalling.

### SOX2 downregulation reconfigures β-CATENIN occupancy at cell-state specific CREs

SOX2 is found with WNT signal transducers at a large number of CREs in CEpiLCs (Koch et al., 2017) and naïve pluripotent ES cells (Boyer et al., 2005) (Cole et al., 2008) (Yi et al., 2011). We reasoned that SOX2 levels may configure distinct transcriptional responses to WNT signalling by altering the binding profile of β-catenin. To investigate this we performed ChIP-seq for SOX2 and β-catenin from naïve ES cells cultured in 2i and CEpiLCs, SOX2-ON and SOX2-OFF cells cultured in FLC medium. We generated consensus peak sets for each factor (Fig 3A) (see methods). This included 76% of SOX2 and 89% of β-catenin peaks identified in an independent study of WT CEpilCS (Koch et al., 2017), plus an additional 110893 SOX2 and 81832 β-catenin peaks (Fig S3A,B).

**Figure 3.**
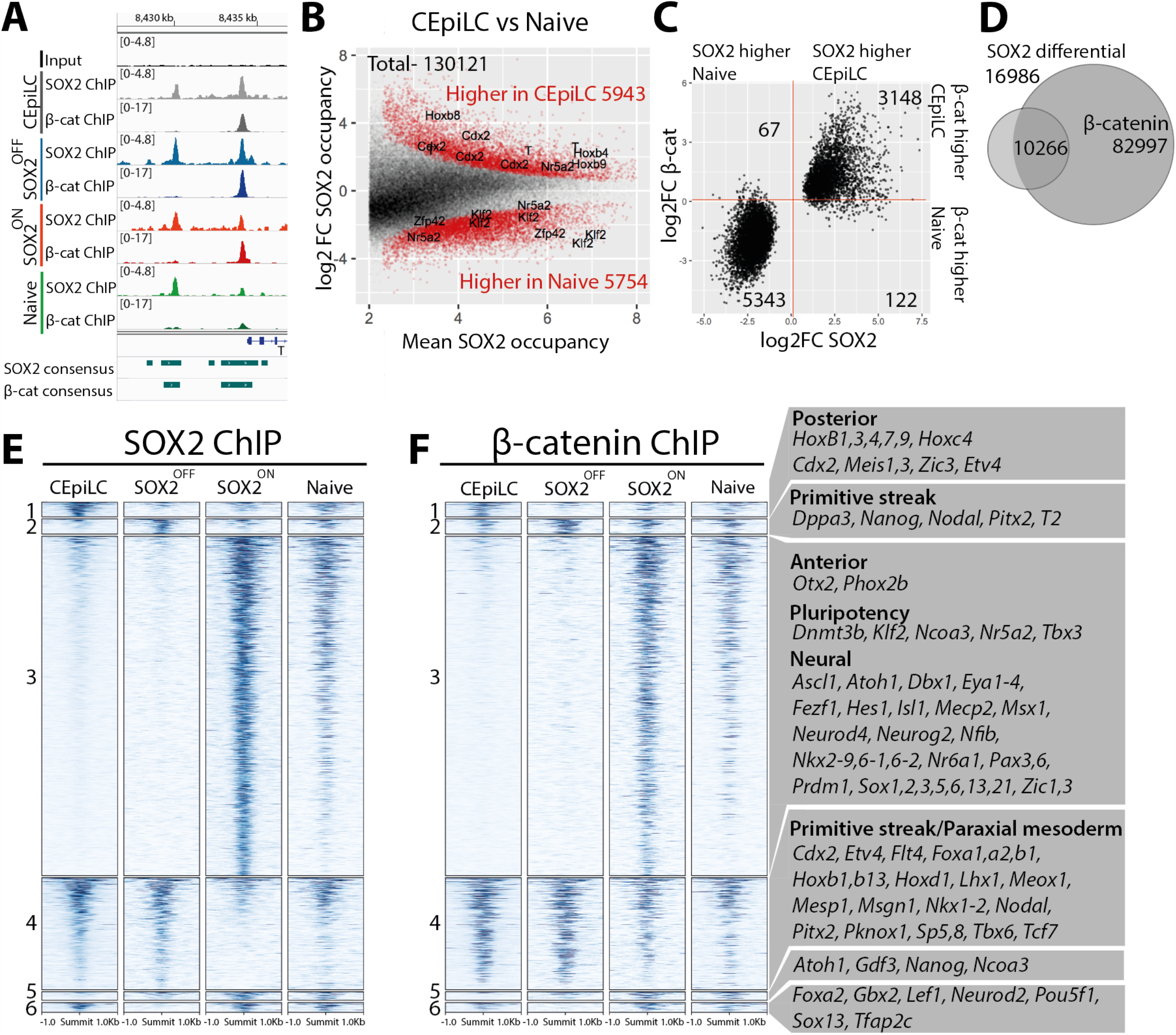
SOX2 downregulation reconfigures β-catenin occupancy. (A) Representative SOX2 and β-catenin ChIP-seq signals used to define high-confidence consensus peaks (see methods). (B) SOX2 occupancy increases at peaks associated with posterior genes and decreases at peaks associated with pluripotency genes during the differentiation of pluripotent ES cells to CEpiLCs. Differentially occupied peaks coloured red are statistically different between conditions. FDR <0.05, n = 3. (C) Peaks differentially occupied by SOX2 between ES cells and CEpiLCs exhibit a correlated increase or reduction of β-catenin occupancy. (D) The majority of peaks differentially occupied by SOX2 (FDR <0.05) across any pairwise comparison between CEpiLC, SOX2 OFF, and SOX2 ON overlap with β-catenin consensus peaks. (E) Cell-type specific SOX2 and (F) β-catenin occupancy at SOX2 differential peaks. Representative nearest genes associated with SOX2 peaks from each cluster are shown. For details of clustering see Methods.

To investigate how the reduction of global SOX2 levels during the transition from pluripotent to CEpiLC progenitors changes SOX2 occupancy genome-wide, we performed differential analysis (Fig 3B). This revealed a surprisingly dynamic pattern of SOX2 binding. SOX2 occupancy was reduced at 5754 sites in CEpiLCs but, unexpectedly, increased at 5943 locations, despite its lower expression levels. Peaks exhibiting higher SOX2 occupancy in naïve ES cells included a set associated with pluripotency genes, whereas peaks exhibiting higher SOX2 occupancy in CEpiLCs were associated with genes characteristic of primitive streak and trunk identity (Fig 3B). Strikingly, β-catenin exhibited a coordinated reconfiguration in its occupancy at sites differentially occupied by SOX2 (Fig 3C). Similarly, the majority of peaks differentially occupied by β-catenin reflected the altered SOX2 occupancy at these sites in CEpiLCs compared to naïve ES cells (5851/5908; 99%) (Fig S3C). This indicated that the changes in SOX2 levels accompanying the transition from pluripotency to CEpiLC might directly reconfigure the transcriptional response to WNT signalling by redistributing β-catenin occupancy.

To test whether changes in SOX2 levels can account for the reconfiguration of SOX2/β-catenin occupancy during the transition from pluripotency to CEpiLC we assayed the effect of experimentally manipulating SOX2 levels in cells cultured under CEpiLC differentiation conditions using the SOX2^TetON^ line. Of the 16986 peaks differentially occupied in response to experimental manipulation of SOX2 levels, 36% were differentially occupied by SOX2 between naïve pluripotent ES cells and CEpiLC. Notably, 60% of SOX2 peaks differentially occupied in response to changes in SOX2 levels overlapped with β-catenin peaks (Fig 3D). To investigate the relationship between SOX2 and β-catenin occupancy we clustered SOX2 occupied CREs to reflect their cell-type specific occupancy (see methods). As observed during the transition from pluripotency to CEpiLC identity, changes in β-catenin occupancy mirrored those of SOX2 (Fig 3E,F, Fig S3D,E). Moreover, both SOX2 and β-catenin occupancy were strikingly similar between both CEpiLC and SOX2 OFF cells (which express low SOX2) (R= 0.84 SOX2, 0.90 β-catenin), and SOX2-ON and naïve progenitors (which express high SOX2) (R= 0.68 SOX2, 0.67 β-catenin). These data provide evidence of a profound genome-wide alteration in the binding site occupancy of β-catenin that is dependent on SOX2.

### SOX2 levels configure TCF/LEF occupancy

SOX factors have been suggested to modulate the association of β-catenin with chromatin via TCF/LEF dependent and independent mechanisms (Mukherjee et al., 2020). Moreover, TCF/LEF factors are differentially expressed between high and low SOX2 expressing cell types (Fig S3F). We therefore performed ChIP-seq with TCF7L1, TCF7L2 and LEF1 in ES cells, CEpiLCs, SOX2 OFF, and SOX2 ON. This revealed that 60% (LEF1), 59% (TCF7L1), and 47% (TCF7L2) of differentially occupied TCF/LEF1 sites overlapped with differentially occupied SOX2 sites (Fig S3G-I). Similar overlaps were observed with differentially occupied β-catenin sites, which showed 69% (LEF1), 65% (TCF7L1), and 63% (TCF7L2) overlap, suggesting occupancy occurs at the same CREs (Fig S3 J-L). Indeed, TCF/LEF factors exhibited a similar pattern of cell-state specific occupancy to SOX2/β-catenin (compare Fig 3E,F, to Fig S3M-O; Fig S3D,E to S3P-R). Taken together these data indicate that the reduction in SOX2 levels during the transition from pluripotency to CEpiLC identity drives the co-ordinated reconfiguration of SOX2/TCF/β-catenin co-occupancy across the genome.

### Redistribution of TCF/B-catenin occupancy establishes the distinct transcriptional response of CEpiLCs

As TCF/β-catenin mediates the transcriptional response to WNT signalling, we asked whether the changes in SOX2/β-catenin binding could explain the distinct gene expression programmes of cells expressing low or high levels of SOX2. For each cluster of differential SOX2 binding peaks, we identified the corresponding sets of differentially expressed neighbouring genes and compared their levels in CEpiLC, SOX2-OFF, and SOX2-ON cells. In line with the known positive effect of β-catenin on transcription, average changes in the expression of genes associated with each cell-state specific cluster were positively correlated with changes in SOX2/β-catenin occupancy (Fig 4).

**Figure 4.**
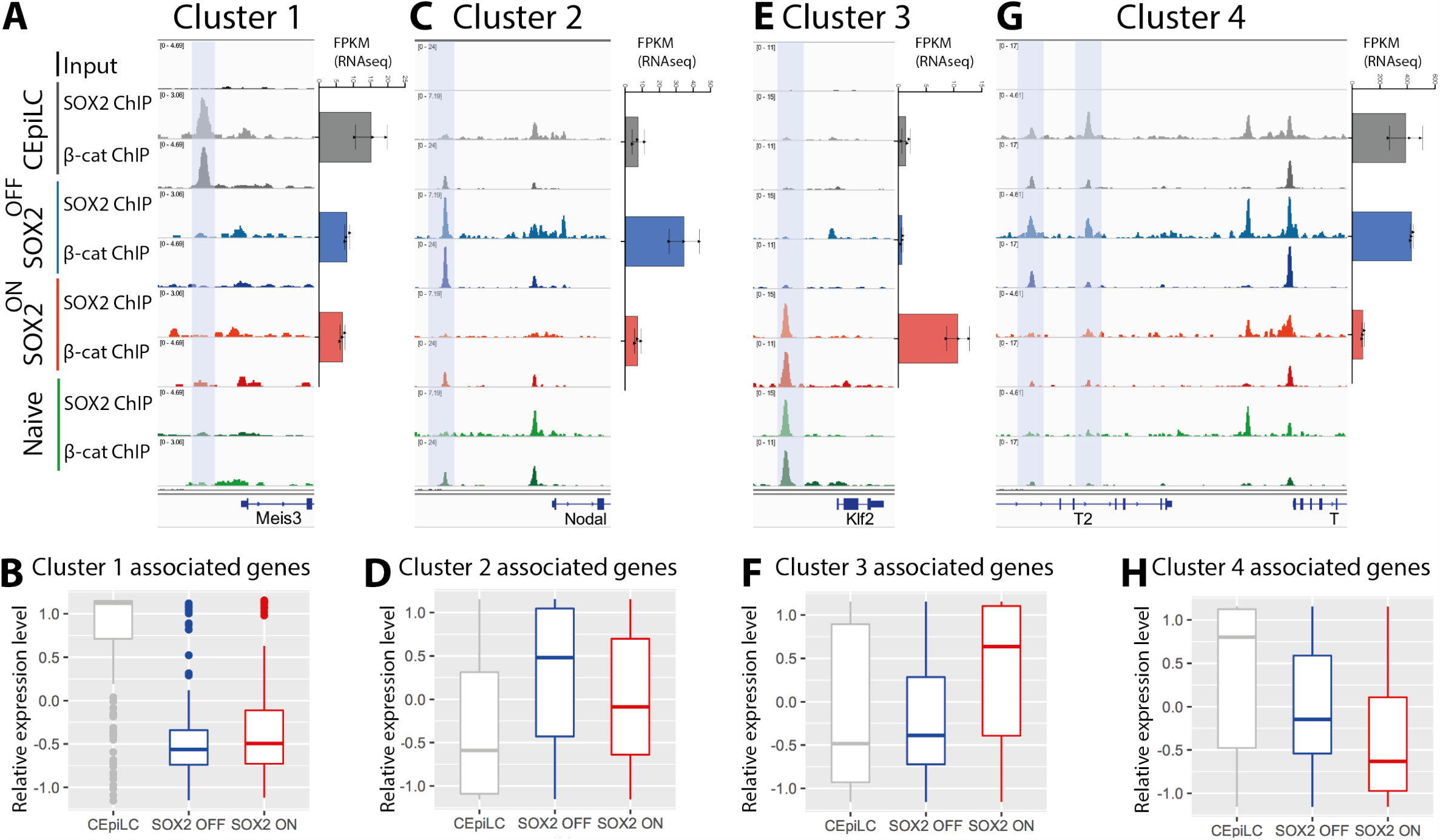
SOX2/β-catenin co-occupancy correlates with cell-type specific gene expression. (A) SOX2/β-catenin co-occupancy at a CRE shaded blue upstream of the transcriptional start site of Meis3 correlates with its cell-type specific expression in CEpiLCs. (B) Differentially expressed genes associated with cluster 1 CREs are most highly expressed in CEpiLCs. (C) SOX2/β-catenin co-occupancy at an upstream of the transcriptional start site of Nodal correlates with its cell-type specific expression in SOX2-OFF cells. (D) Differentially expressed genes associated with cluster 2 CREs are most highly expressed in SOX2-OFF. (E) SOX2/β-catenin co-occupancy at a CRE upstream of the pluripotency marker *Klf2* correlates with its cell-type specific expression in SOX2-ON cells. (F) Differentially expressed genes associated with cluster 3 CREs are most highly expressed in SOX2-ON. (G) Reduced SOX2/β-catenin co-occupancy at multiple CREs upstream of the transcriptional start site of *T*/*Bra* in SOX2 ON and Naïve ES cells correlate with the absence of T/Bra expression in those cell-states. (H) Differentially expressed genes associated with cluster 4 CREs are expressed at lowest levels in SOX2-ON. Differential expression criteria = FDR < 0.05.

Genes in cluster 1, which are specifically occupied by SOX2/β-catenin in CEpiLCs (Fig 4A, Fig 3E,F) and reduced in expression in both SOX2-ON and SOX2-OFF progenitors (Fig 4-A,B), were enriched for biological processes related to anterior-posterior patterning (Fig S4A). This included known posterior determinants *Cdx2* and *HoxB1-9*, Hox co-factors *Meis1* and *Meis3*, and posterior markers *Etv4* and *Zic3* (Fig 3F). Genes in cluster 2 were occupied by SOX2/β-catenin most highly in SOX2-OFF cells (Fig 4C, Fig 3E,F) and included early primitive streak and mesendodermal genes *Nodal, Pitx2*, and *Nanog* (Fig 3F). These showed greatest average expression in SOX2-OFF (Fig 4C,D). Cluster 3 genes showed greatest SOX2/β-catenin occupancy and expression in SOX2-ON cells (Fig 4E, Fig 3E,F) and comprised genes associated with pluripotency and anterior neural identity (Fig 3F, Fig S4B). Genes in cluster 4, which are occupied by SOX2/β-catenin most highly in WT CEpiLCs and SOX2-OFF cells (Fig 4G, Fig 3E,F) were enriched for genes expressed in caudal epiblast (*Sp5/8, Nkx1-2, Cdx2*), primitive streak (*Foxa1/2, Pitx2, Nodal, Lhx1, T/Bra*) and mesoderm (*Meox1, Mesp1, Msgn1, Tbx6*) (Fig 3F). Similar to cluster 1 and 2, depletion of SOX2/β-catenin occupancy at cluster 4 CREs in SOX2-ON cells was correlated with reduced expression compared to both WT EpiLCs and SOX2-OFF cells (Fig 4H). Thus, differential SOX2/β-catenin occupancy driven by changes in SOX2 levels correlate with cell-state specific gene expression patterns and points to a positive role for SOX2 in promoting WNT-dependent gene activation by β-catenin in caudal epiblast, primitive streak, and pluripotent progenitors.

### High SOX2 levels direct TCF/β-catenin occupancy to pluripotency and neural genes

Differentially expressed genes associated with SOX2/β-catenin occupancy in SOX2-ON in CEpiLC conditions and naïve progenitors (cluster 3 and 5), included a number of pluripotency factors (*Dnmt3b, Klf2, Klf5, Ncoa3,Nr5a2, Tbx3*) as well as genes involved in nervous system development (Fig 3F, S4-B). Neural genes were expressed at comparatively low levels (Fig S4C), consistent with the priming of these genes in embryonic stem cells (Bergsland et al., 2011). Thus the establishment of a naïve-like SOX2/TCF/LEF configuration appears to underlie the re-expression of pluripotent factors in SOX2-ON cells stimulated with WNT agonist, and repression of the post-implantation epiblast WNT response. This supports the idea that a reduction in SOX2 levels is necessary to allow reconfiguration of SOX2 and β-catenin to ensure a caudal epiblast gene regulatory program in response to WNT activity.

### High SOX2 levels increase chromatin accessibility at pluripotency and neural regulatory elements

SOX2 has been proposed to act as a pioneer factor that can promote the occupancy of TFs at otherwise inaccessible chromatin (Soufi et al., 2015) (Dodonova et al., 2020) (Soufi et al., 2012) (Malik et al., 2019), suggesting that SOX2 may direct cell-state specific TCF/β-catenin binding by altering chromatin accessibility. To test this possibility, we performed ATAC-seq analysis and quantified accessibility across regions exhibiting cell-state specific differential SOX2 occupancy. This analysis revealed distinct relationships between chromatin accessibility and SOX2 occupancy between cell-states. Cluster 3 CREs, which are associated with pluripotency and neural progenitor genes, and occupied by SOX2 in SOX2-ON and pluripotent progenitors, were only accessible in cell-states with high SOX2 (Fig 5A,B; S5A). By contrast, CREs in cluster 1, which showed SOX2 binding in CEpiLC, were accessible to a comparable degree in all cell-states (Fig 5A, S5A). Cluster 2 (SOX2 OFF-specific/early streak) and cluster 4 (primitive streak/paraxial mesoderm), exhibited a more complex pattern of accessibility, with comparable average accessibility in SOX2-OFF, SOX2-ON, and pluripotent progenitors, but less accessibility in CEpiLCs (Fig 5A, S5A). Thus, whereas changes in chromatin accessibility may explain the specific occupancy of SOX2/TCF/β-catenin complexes in SOX2-ON and pluripotent cells, which express high levels of SOX2, they do not explain the cell-state specific occupancy at CEpiLC-specific cluster 1, SOX2-OFF specific cluster 2, or primitive streak/paraxial mesoderm cluster 4, in low SOX2 expressing progenitors.

**Figure 5.**
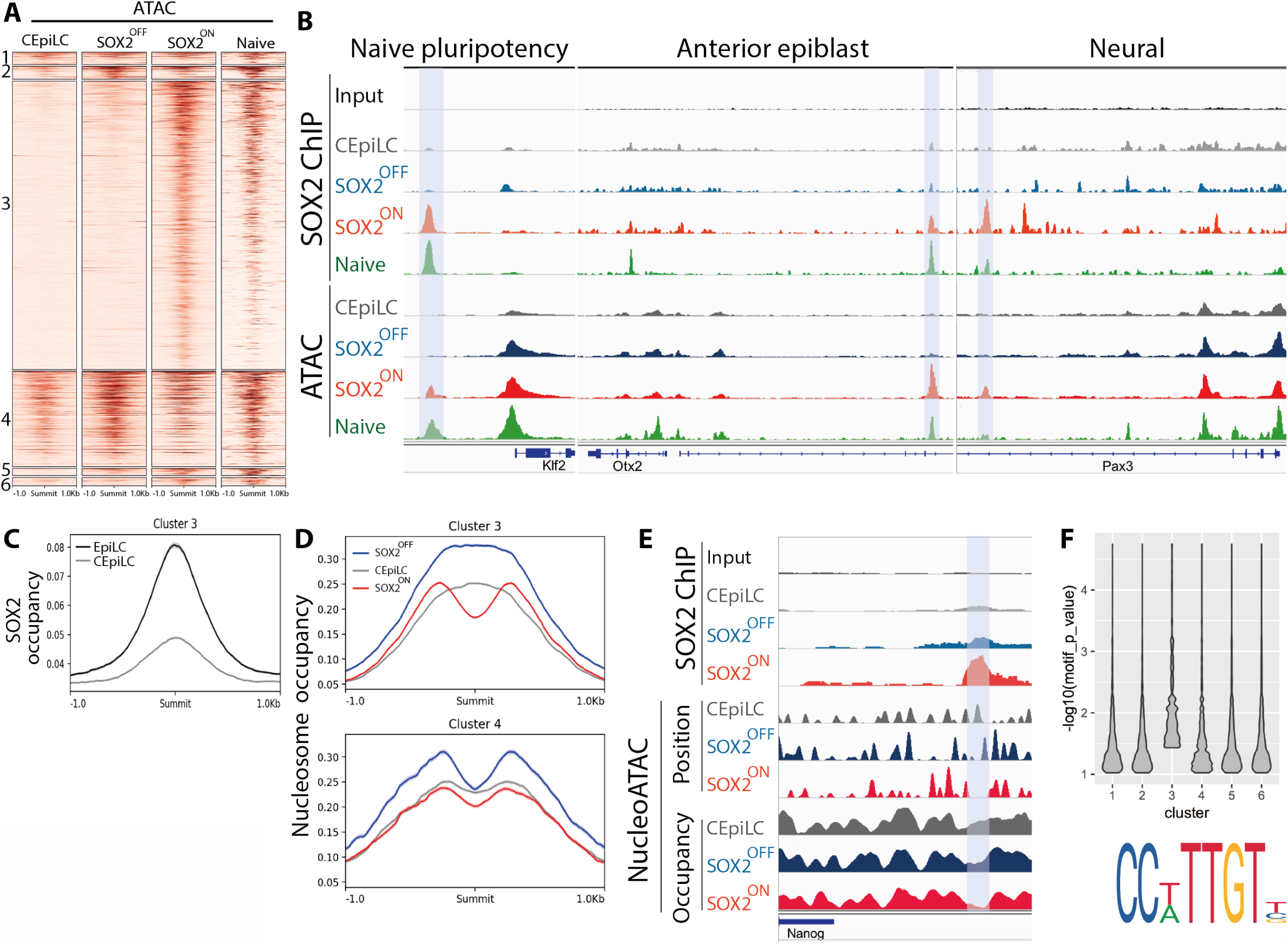
SOX2 promotes chromatin accessibility at high-affinity sites. (A) Cluster 3 CREs classified in Fig 3E exhibit ATAC-seq accessibility specifically in high SOX2 expressing cells. Naïve ATAC-seq data reanalysed from (Kearns et al., 2015). (B) Correlated SOX2 occupancy and ATAC-seq accessibility at representative cluster 3 CREs (shaded blue) associated with genes expressed in high SOX2 expressing cell types. (C) Cluster 3 peaks exhibit greater ATAC-seq accessibility in EpiLCs than in CEpiLCs (reanalysed from (Metzis et al., 2018)). (D) Average nucleosome occupancy at cluster 3 CREs is reduced at SOX2 peak centres in SOX2-ON, but relatively unchanging between cell types at cluster 4 peaks. (E) Negative-correlation between SOX2 occupancy, nucleosome position, and nucleosome occupancy at a representative CRE (shaded blue). (F) SOX2 binding motifs at cluster 3 peaks most closely resemble the consensus sequence shown.

Accessibility at SOX2 occupied sites in cluster 3 might be a direct consequence of elevated SOX2 or an indirect consequence of the WNT-dependent naïve pluripotent cell-state. To distinguish between these two possibilities we compared accessibility at these sites between WT CEpiLCs (low SOX2 levels) and EpiLCs, which have high levels of SOX2 but do not express pluripotency genes (see Fig 2G). This revealed that accessibility at cluster 3 peaks was on average higher in EpiLCs than in CEpiLCs (Fig 5C, S5B), and correlated with SOX2 occupancy at both pluripotent and neural gene associated CREs (Fig S5C). This supports the idea that high SOX2 levels maintain accessibility at cluster 3 sites independently of additional pluripotency factors.

We explored whether the increased ATAC-seq signal at cluster 3 sites in SOX2-ON was driven directly by SOX2 occupancy by analysing the nucleosome landscape at sites of SOX2 binding. NucleoATAC analysis (Schep et al., 2015) revealed that average nucleosome occupancy at the centre of cluster 3 SOX2 binding peaks was markedly depleted in high SOX2 expressing SOX2-ON cells compared to low SOX2 expressing SOX-OFF cells and WT CEpiLCs (Fig 5D). By contrast, the average nucleosome density at SOX2 peak centres in cluster 4 peaks was largely independent of SOX2 levels, consistent with the largely unchanging chromatin accessibility within these peak clusters across cell-states (Fig 5A, S5A). As chromatin accessibility at cluster 3 peaks is correlated with reduced nucleosome occupancy specifically at SOX2 binding sites (Fig 5E) we conclude that SOX2 binding directly drives nucleosome eviction, rather than being a secondary consequence of increased neighbouring accessibility.

We hypothesised that the distinct relationship between SOX2 levels and nucleosome occupancy at cluster 3 peaks may reflect the underlying sequence of SOX2 binding sites. Putative SOX2 binding sites located at the centres of cluster 3 peaks more closely matched the consensus motif than those in the other clusters (Fig 5F), indicating that they are of higher affinity than those found in other peaks. What then explains the change in SOX2 binding in CEpiLC cells?

### SOX2/TCF/LEF co-occupy low affinity SOX2 sites with cell-state specific factors in primitive streak progenitors and CEpiLCs

As cell-type specific TF binding at low affinity sites is associated with cooperative co-factor binding (Slattery et al., 2011) (Crocker et al., 2015), we reasoned that SOX2 occupancy at constitutively accessible sites might require additional cell-state specific co-factors. To investigate this, we performed motif enrichment analysis (Fig 6A). Cluster 1 peaks, which exhibit SOX2 occupancy specifically in CEpiLC conditions, were enriched for CDX/HOX motifs, indicating that CEpiLC specific SOX2/TCF/β-catenin occupancy may be directed by binding with CDX factors expressed specifically in CEpiLCs. Similarly, cluster 4 sites, which are occupied by SOX2/TCF/β-catenin in both CEpiLCs and SOX-OFF primitive streak progenitors, were enriched for motifs for T/BRA, in addition to OCT, OCT/SOX/TCF/NANOG, and TCF/LEF motifs, as were SOX-OFF specific cluster 2 sites. Notably, cluster 2 sites were also enriched for the Nodal signalling mediator FOXH1, indicating that elevated Nodal signalling may contribute to the regulation of the distinct early primitive streak WNT response in SOX-OFF cells.

**Figure 6.**
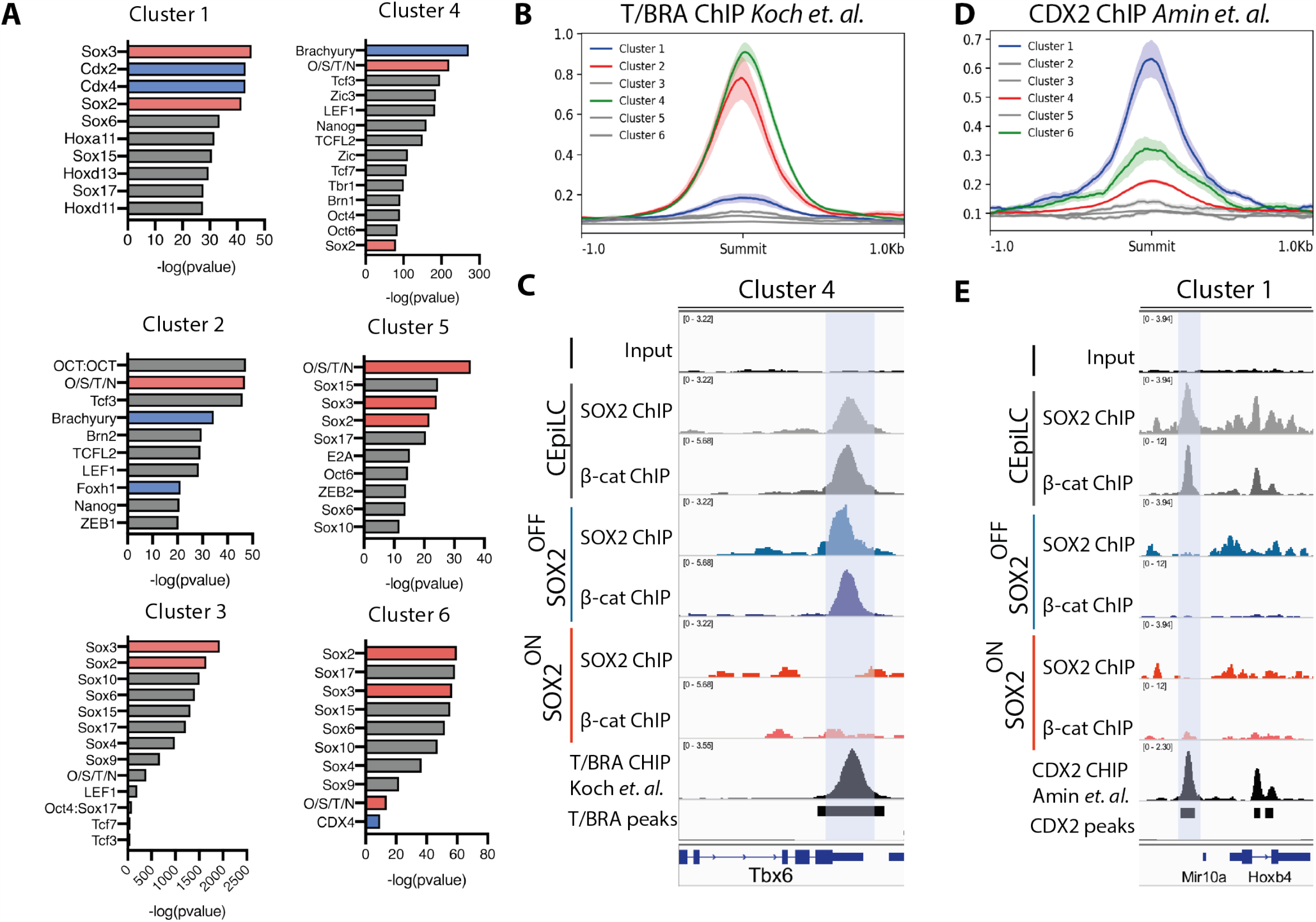
SOX2 associates with cell-type specific factors at low-affinity sites. (A) Cluster 1-6 CREs are enriched for distinct sets of TF binding motifs. (B) Cluster 2 and 4 CREs are enriched for T/Bra occupancy. T/BRA ChIP-seq reanalysed from (Koch et al., 2017). (C) Correlated SOX2, β-catenin and T/Bra occupancy at a representative cluster 4 CRE (shaded blue) at the*Tbx6* locus. (D) Cluster 1 CREs are enriched for CDX2 occupancy. CDX2 ChIP-seq reanalysed from (Amin et al., 2016). (E) Correlated SOX2, β-catenin and CDX2 occupancy at a representative cluster 1 CRE (shaded blue) at the HoxB locus. O/S/T/N = OCT4/SOX2/TCF7L1/NANOG. All bars in metaplots = sem.

As T/BRA is expressed in both CEpiLCs and SOX2-OFF progenitors but repressed in SOX2-ON progenitors, T/BRA may mediate SOX2/TCF/LEF occupancy at cluster 2 and 4 sites. Consistent with T/BRA facilitating low-affinity SOX2 binding, analysis of ChIP-seq data from WT CEpiLCs indicated that T/BRA was enriched, along with SOX2 and β-catenin, at both cluster 2 and cluster 4 sites (Fig 6B,C; Fig S6) and was most strongly associated with CREs with low levels of SOX2 occupancy. Similarly, CDX2 ChIP-seq data from CEpiLCs confirmed that the enrichment for CDX2 motifs within cluster 1 was mirrored by an enrichment for CDX2 co-occupancy with SOX2 and β-catenin (Fig 6D,E; Fig S6.

### CDX2 enhancer activity is reduced in high and low SOX2 expressing progenitors

To test directly the role of SOX2 in regulating CEpiLC specific gene expression we focused on CDX2. CDX2 expression is constrained to a specific range of SOX2 levels (Fig 2A, Fig S7A) and likely contributes to both the distinct SOX2/β-catenin occupancy and WNT response of CEpiLCs. Inspection of the CDX2 locus identified a putative regulatory element within the CDX2 intron overlapping a previously identified enhancer (Gaunt et al., 2005) (Wang and Shashikant, 2007) which exhibited a cell-type specific pattern of SOX2/β-catenin co-occupancy that correlated with SOX2 levels (Fig 7A). To test the activity of SOX2 at this element we generated fluorescent reporter lines harbouring the intronic sequence (Fig 7-B). Both CDX2 expression and reporter activity were higher in CEpiLC cells cultured in FLC medium compared to pluripotent cells, which express high levels of SOX2, and activin-induced early primitive streak cells (Fig S7B, C) that express little if any of SOX2 or its functionally equivalent homolog *Sox3* (Fig 7C-E, Fig S7D). We constructed a variant of the reporter in which the eight identifiable SOX2 binding sites were scrambled (Sox2del) (Fig 7B). Activity of the Sox2del reporter was substantially reduced in CEpiLC (Fig 7F, Fig S7E,F) consistent with the idea that SOX2 occupancy promotes the induction of CDX2 by WNT signalling.

**Figure 7.**
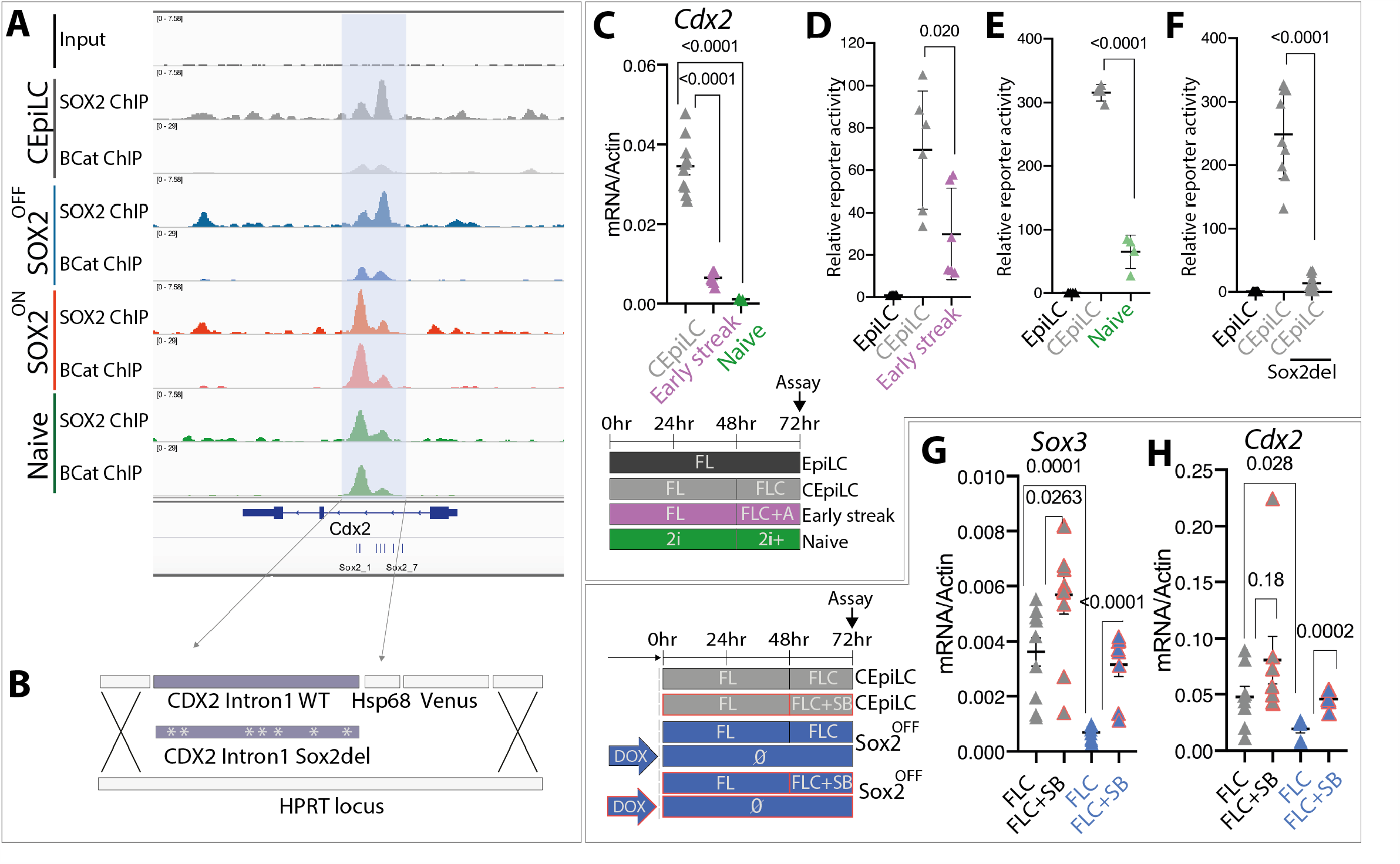
Cdx2 induction requires low-level SOX2/SOX3 expression. (A) SOX2 and β-catenin co-occupy a CRE within intron 1 of *Cdx2* (shaded blue). (B) Wild-type and modified sequence (Sox2del) from the *Cdx2* intronic CRE was cloned into a fluorescent reporter construct. (C) *Cdx2* expression is reduced in pluripotent and activin-induced early streak progenitors. CHIR concentration was increased to 5uM in 2i+ conditions. (D) The activity of the *Cdx2* CRE reporter construct is reduced in early streak and (E) pluripotent progenitors relative to caudal epiblast progenitors (CEpiLC). (F) Deletion of SOX2 binding sites (Sox2del) reduces the activity of the *Cdx2* CRE reporter in CEpiLCs. (G) Inhibition of nodal signalling elevates *Sox3*, and (H) *Cdx2* expression in SOX2-OFF progenitors. Individual biological replicates are plotted, and p-values are show.

SOX2 and CDX2 are repressed by Nodal signaling in early primitive streak progenitors (Mendjan et al., 2014) (Teo et al., 2011). As Nodal expression is elevated in SOX2-OFF primitive streak progenitors (see Fig 3D) this suggests a mechanism by which the reduction in SOX2 levels establishes a feedback loop via Nodal signalling that further inhibits the CEpiLC-specific WNT response. To test this we inhibited Nodal signalling in SOX-OFF cells concurrently with WNT pathway activation. This led to the inhibition of both the general primitive streak marker T/BRA, expression of early primitive streak markers *Eomes, Mixl1*, and *Nanog* (Fig S7G), upregulation of *Sox3* (Fig 7G), and to a rescue of *Cdx2* expression (Fig 7H). We conclude that the presence of moderate levels of SOX2 in CEpiLCs promotes posterior identity by both positively regulating CDX2 expression and restraining the induction of early primitive streak identity by Nodal.

## Discussion

Here we show that the level of SOX2 expression determines its genome-wide occupancy and this underpins distinct WNT-driven transcriptional programmes at sequential stages of pluripotent stem cell differentiation. We found that β-catenin frequently co-occupies genomic sites with SOX2 and that changes in the genomic location of SOX2 led to coordinated changes in chromatin binding of β-catenin. During the transition from pluripotency to caudal epiblast identity a reduction in global SOX2 levels resulted in a reduction of SOX2 occupancy at a set of CREs and this was accompanied by a corresponding reduction in β-catenin occupancy. Many of these CREs were associated with genes expressed in pluripotent epiblast or neural ectoderm progenitors, cell types that require high levels of SOX2 expression to maintain their identity (Avilion et al., 2003) (Wang et al., 2012) (Bergsland et al., 2011) (Bylund et al., 2003). Surprisingly, we observed that the reduction in global SOX2 levels also led to an increase in SOX2 and β-catenin co-occupancy at a distinct set of CREs. These were associated with WNT-responsive genes expressed in caudal epiblast progenitors, many of which are responsible for posterior patterning and mesoderm differentiation. This depends on the precise levels of SOX2. Artificially increasing or decreasing SOX2 expression redistributed SOX2/β-catenin, and prevented the transition to a CLE identity. This demonstrates a role for SOX2 levels in configuring the WNT response of epiblast progenitors and shaping the transcriptional changes accompanying the differentiation of pluripotent cells to caudal lateral epiblast.

Following SOX2 downregulation, the loss of SOX2 binding at CREs associated with neural and pluripotency genes correlated with a loss in chromatin accessibility. This is consistent with the known role of SOX2 as a pioneer-factor and its ability to bind and open inaccessible CREs (Soufi et al., 2015) (Dodonova et al., 2020) (Soufi et al., 2012) (Malik et al., 2019). In vivo, epiblast progenitors share a common epigenetic configuration and chromatin accessibility profile with ectodermal lineages (Argelaguet et al., 2019), consistent with the tendency of ES and epiblast stem cells to differentiate to neural cells (Tropepe et al., 2001). The occupancy of these CREs by SOX2 suggests that the neural CREs are maintained in an accessible primed state, ready for expression following exit from pluripotency (Rada-Iglesias et al., 2011) (Bergsland et al., 2011). A corollary of this is that decreased SOX2 expression, which accompanies differentiation of pluripotent cells to fates other than neural, is responsible for the closure and epigenetic silencing of CREs associated with neural identity (Argelaguet et al., 2019).

The decrease in SOX2 levels resulted in a repositioning of SOX2 to CREs associated with genes involved in AP patterning and mesoderm induction. Surprisingly, despite the lower levels of SOX2, these CREs contained lower affinity SOX2 binding sites than the CREs bound by SOX2 in cells types with high SOX2 expression levels. Moreover, these CREs were accessible in pluripotent conditions as well as in caudal epiblast. Cofactor mediated recruitment to low affinity sites has been implicated in cell type specific CRE activity and gene expression (Slattery et al., 2011) (Crocker et al., 2015) (Farley et al., 2015). For example, the recruitment of the endoderm pioneering factor FOXA2 to regulatory elements with lower affinity binding sites depends on lineage-specific TFs (Geusz et al., 2020). For SOX2, we found evidence of the involvement of CDX2 and T/BRA in directing binding to low affinity sites. Taken together, these observations suggest that SOX2 adopts different modes of chromatin interaction and CRE selection depending on its level of expression. This resolves a paradox. Despite its pioneering activity and ability to bind and activate condensed chromatin, the distribution of SOX2 occupancy on chromatin differs between cell types. Our data provide evidence that SOX2 acts as a pioneer factor in pluripotent cells when expressed at high levels, but collaborates with other transcription factors to select lower affinity binding sites when expressed at lower levels. This provides a causal explanation for how modulating TF expression levels can regulate distinct gene expression programmes.

There was a positive correlation between SOX2 binding and the activation of WNT responsive genes in caudal lateral epiblast cells. Consistent with this, using a CRE from CDX2 as a model, we found that SOX2 occupancy is required for CDX2 activation by WNT signalling, providing direct evidence for an activator role of SOX2 in the regulation of β-catenin target genes. This suggests a self-reinforcing mechanism regulating cell-type specificity of WNT signalling. Downregulation of SOX2 leads to the dissolution of the pluripotent chromatin state and transcriptional programme. This eases repression on caudal epiblast specific WNT target genes such as T/BRA and CDX2. Consequently, SOX2 and TCF/β-catenin are recruited to CREs associated with CDX and T/BRA target genes, inducing the expression of gene expression programmes characteristic of posterior identity and primitive streak/paraxial mesoderm differentiation. Then, as SOX2 levels are further reduced during the differentiation of caudal epiblast progenitors to mesoderm progenitors (Javali et al., 2017) (Wymeersch et al., 2016), CDX and BRA expression decreases.

The chromatin locations that are differentially occupied by β-catenin are enriched with binding sites for TCF/LEF, interacting partners of β-catenin (Steinhart and Angers, 2018) (Cadigan and Waterman, 2012). SOX family factors have been demonstrated to interact with TCF/LEF/β-catenin complexes via a number of mechanisms. SOX17 physically interacts with both β-catenin and TFC/LEF factors and recruits β-catenin to chromatin both in collaboration with and independently of TCF/LEF factors in order to regulate mesendodermal gene expression (Mukherjee et al., 2020). Moreover, SOX factors have been shown to either promote the stability or degradation of β-catenin (Akiyama et al., 2004). Our data are consistent with the idea that SOX2 primarily functions to direct the chromatin binding location of TCF/LEF/β-catenin complexes. TCF7L1 co-occupies CREs with SOX2 at OCT/SOX/NANOG/TCF sites associated with pluripotency and neural genes, and promotes pluripotency gene expression in the presence of WNT in ES cells. Moreover, forced expression of high levels of SOX2 in epiblast-like cells was sufficient to drive TCF7L1 occupancy at CREs associated with both pluripotency and neural genes, and to reactivate pluripotent gene expression in the presence of WNT signalling. In caudal epiblast progenitors, which preferentially express LEF1 over TCF7L1, SOX2 and LEF1 co-occupancy is elevated at genes associated with AP patterning and paraxial mesoderm differentiation. We also found that the occupancy of TCF7L2, which is not differentially expressed between high and low SOX2 progenitors, mirrors the changes in SOX2 occupancy, suggesting that SOX2 promotes the recruitment of specific TCF/LEF factors to distinct CREs. As TCF/LEF factors have been shown to be largely redundant (Gerner-Mauro et al., 2020) (Moreira et al., 2017), the significance of this specificity is unclear. We note that our analysis does not distinguish whether direct physical interaction between SOX2 and β-catenin mediates recruitment to TCF/LEF complexes. Moreover, although we cannot rule out the possibility that TCF/LEF act as pioneer factors to drive SOX2 occupancy in high SOX2 expressing progenitors, we demonstrate the reconfiguration of SOX2/TCF/LEF/β-catenin co-occupancy is a direct consequence of changes in SOX2 levels.

In contrast to the primed pluripotent epiblast, naïve progenitors maintain high SOX2 levels in the presence of WNT signalling. How is the molecular identity of primed pluripotent progenitors altered such that WNT signalling inhibits SOX2 expression in these cells? The transition to primed pluripotency is acquired in response to FGF pathway activation (Kunath et al., 2007), hence FGF target genes represent candidates for mediating the transition. The FGF pathway mediator ETV5 has been shown to drive the naïve to primed pluripotency transition in the presence of FGF, leading to upregulation of ETV4 (Kalkan et al., 2019). It will be interesting to determine whether ETV factors play a direct role in repressing SOX2 in caudal epiblast progenitors, or whether this is mediated by ETV-dependent WNT target genes. Moreover, the Hippo pathway mediators YAP/TAZ have been shown both to repress SOX2 in early embryos and are regulated by WNT signalling (Azzolin et al., 2014)(Azzolin et al., 2012)(Frum et al., 2019), suggesting another possible mechanism to downregulate of SOX2.

An interaction between Nodal and WNT signalling has previously been shown to control the differentiation of early and late streak derived mesoderm lineages through a mechanism involving reciprocal antagonism between CDX2 and Nanog (Mendjan et al., 2014). Our data highlight a central role for SOX2 in this switch. Rapid downregulation of SOX2 in early streak progenitors leads to the loss of SOX2 input on CDX2 and the redistribution of SOX2/β-catenin to CREs associated with early streak determinants, including NANOG, PITX2 and Nodal that are enriched for NANOG, T/BRA and FOXH1 binding sites. Moreover, the upregulation of Nodal expression in response to rapid SOX2 downregulation establishes a reinforcing feedback loop by further repressing SOX2, SOX3 and CDX factors and promoting expression of early streak determinants such as NANOG, EOMES and MIXL1. By contrast, in late streak progenitors, SOX2 perdurance leads to the establishment of a CDX2 autoregulatory loop (Xu et al., 1999) while SOX2 β-catenin occupancy is reduced at *Nanog* and *Nodal* regulatory elements. Coupled to the mutual repression between CDX2 and NANOG (Mendjan et al., 2014), the dynamics of SOX2 expression in primitive streak/caudal epiblast progenitors tips the balance between either of two self-reinforcing early or late mesoderm progenitor specifying programs. In addition to repressing CDX2 expression in early mesoderm progenitors, NANOG represses CDX2 in pluripotent epiblast/ES cells (Mendjan et al., 2014) (Chen et al., 2009). NANOG is induced by WNT at both low and high levels of SOX2, with SOX2/β-catenin exhibiting specific patterns of occupancy at the *Nanog* locus depending on SOX2 levels (Fig 3F, Fig 5E). CDX2 is induced in caudal epiblast progenitors expressing intermediate levels of SOX2 at which NANOG is not expressed, indicating that the regulation of NANOG by SOX2 may represent an additional component of the regulatory mechanism that establishes competence for CDX2 expression.

A consequence of the mechanism that establishes the primary body axis is that anterior and posterior structures derive from distinct epiblast progenitor pools (Metzis et al., 2018). Anterior tissues, including the forebrain and heart, are established early in embryonic development from pluripotent epiblast progenitors that co-express OCT4, Nanog, and high levels of SOX2 (Iwafuchi-Doi et al., 2012) (Hart et al., 2004)(Mesnard et al., 2006)(Morgani et al., 2017) (Osorno et al., 2012) (Wood and Episkopou, 1999)(Tam et al., 1997) (Mulas et al., 2018) (Mendjan et al., 2014). By contrast, caudal epiblast progenitors retain low levels of SOX2 expression and this is required to assign trunk identity to both the mesoderm and spinal cord by establishing CDX/HOX expression in response to WNT-signalling. We propose that the potency of caudal epiblast progenitors is a consequence of the low SOX2 levels. This contrasts with earlier epiblast, in which pluripotency is maintained by balancing the repression and activation of determinants of the principal lineages by antagonism between SOX2, OCT4 and NANOG (Thomson et al., 2011) (Teo et al., 2011) (Mulas et al., 2018). As CREs associated with neural genes require high SOX2 to maintain accessibility, neural differentiation is restrained in caudal epiblast progenitors independently of inhibitory activity of OCT4/NANOG. By contrast, the initiation of mesoderm differentiation is controlled by regulation of T/BRA by WNT/Nodal signalling, independently of chromatin remodelling.

A division between the ontogeny of head and trunk tissue is also apparent in arthropods. Reminiscent of the caudal lateral epiblast, homologs of SOX2 and CDX2 are coexpressed in a posterior progenitor pool that fuels WNT dependent axis elongation in arthropods. Moreover, the SOX orthologs have been shown to participate in the assignment of posterior identity within this progenitor population (Copf et al., 2004) (Clark and Peel, 2018)(Bonatto Paese, 2018). Taken together, therefore, the data raise the possibility that a collaboration between WNT signalling and SOX2 in the regulation of CDX factors is an evolutionarily conserved mechanism that establishes the primary bilaterian axis and allocates cells to the body’s trunk tissues.

## Acknowledgements

We are grateful to Tom Frith, Kenzo Ivanovitch and Manuela Melchionda for experimental support and critical feedback during manuscript preparation. We also thank Andreas Sagner and other members of the lab for their generosity sharing insight, expertise and reagents, and the Crick Science Technology Platforms in particular the Advanced Sequencing Facility, Flow Cytometry Facility, and the Bioinformatics and Biostatistics group.

## Author Contributions

R.B. and J.B. conceived the project, interpreted the data, and wrote the manuscript with input from all authors. R.B. designed and performed experiments and data analysis. H.P. performed bioinformatic analysis. T.W. generated reagents. M.G shared reagents, protocols, and unpublished data. V.M. and J.D. provided advice and assistance with ATAC experiments

## Figure legends

**Supplementary Figure relating to Figure 1.**
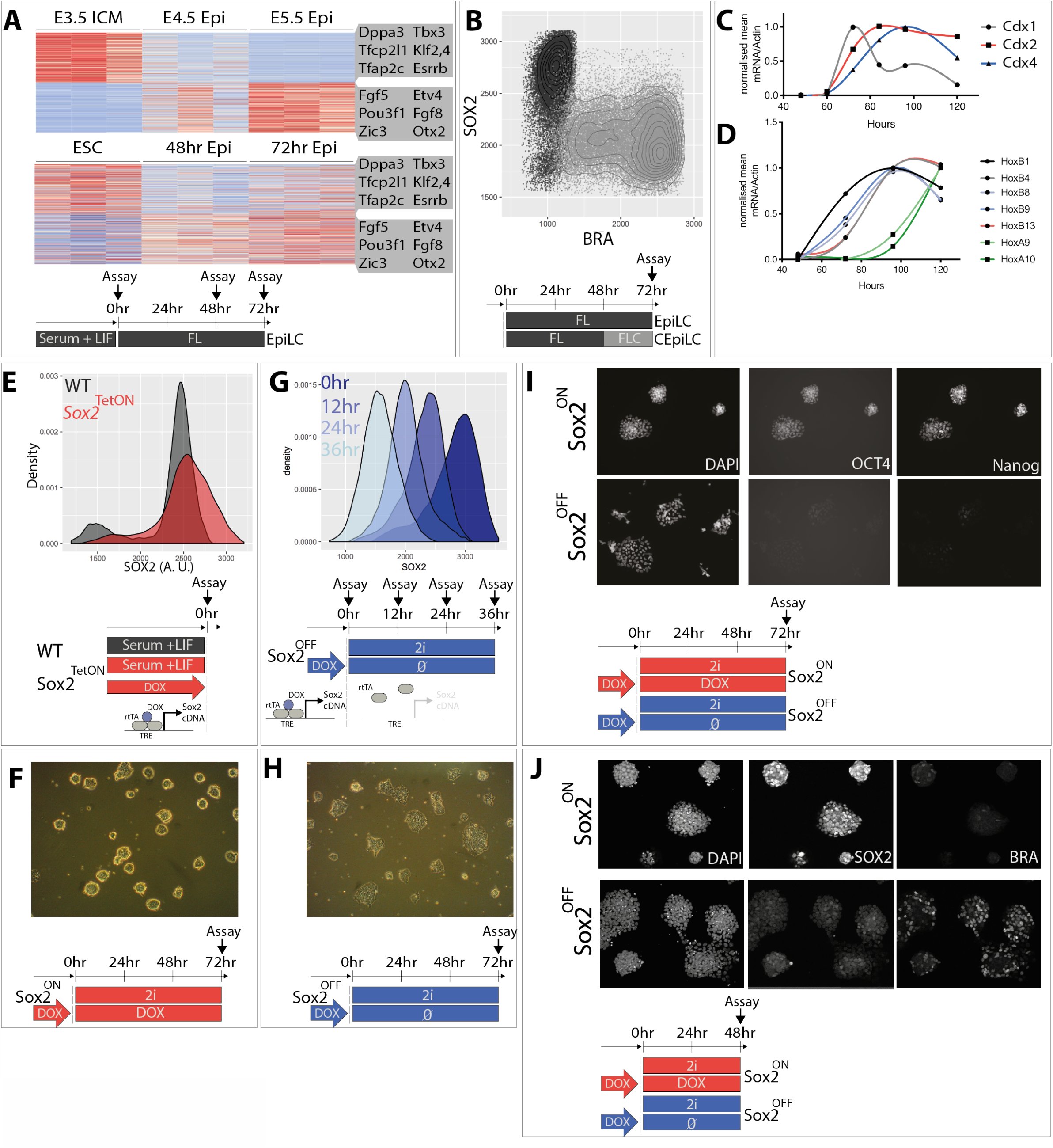
SOX2 maintains pluripotency of SOX2-TetON ES cells. (A) Epiblast-like cells differentiated in FL medium recapitulate in vivo gene-expression dynamics (Boroviak et al., 2015). Illustrative genes for each cluster are highlighted. See methods for details. (B) Culture in FLC medium reduces SOX2 levels and induces T/BRA. (C) FLC medium induces caudal epiblast markers *Cdx1*,2,4, and (D) posterior Hox genes. (E) SOX2-TetON cultured in the presence of doxycycline (SOX2-ON) express SOX2 at comparable levels to pluripotent stem cells, and (F) maintain an undifferentiated morphology in ‘2i’ medium. (G) SOX2 levels are progressively reduced following removal of Dox from SOX2-TetON (SOX2-OFF). SOX2-OFF cultured in ‘2i’ medium lose their undifferentiated morphology (H) and expression of pluripotency markers OCT4 and NANOG (I), and induce the primitive streak marker T/BRA in the absence of SOX2 expression (J).

**Supplementary Figure relating to Figure 2.**
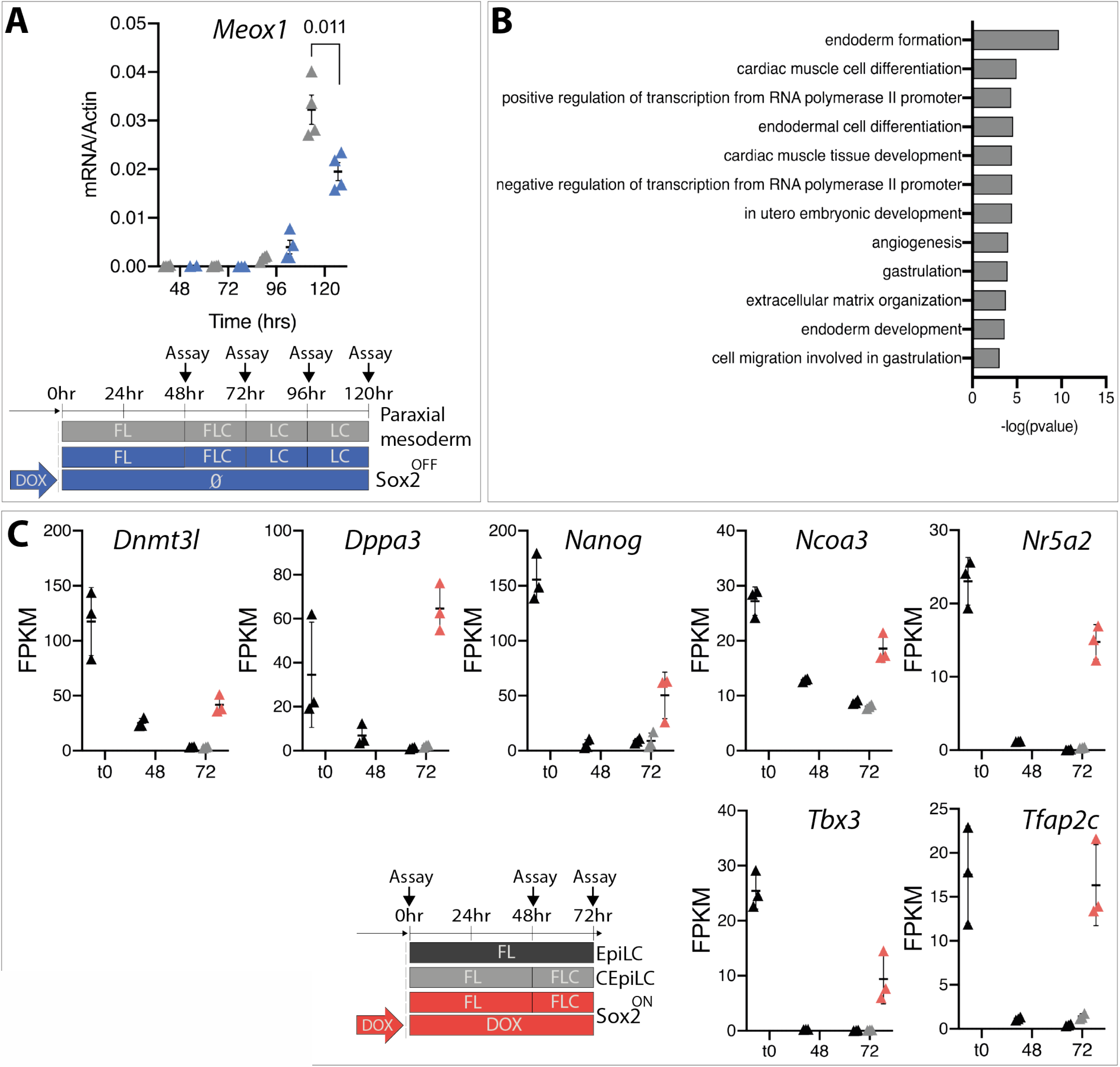
SOX2-OFF and SOX2-ON adopt distinct identities in response to WNT signalling. (A) Somitic mesoderm marker (*Meox1*) expression is reduced in SOX2-OFF compared to CEpiLCs. (B) GO analysis reveals that genes specifically upregulated in SOX2-OFF compared to EpiLC and CEpiLC (from clustering in Fig 2C) are enriched for biological processes indicative of an early/anterior streak identity. (C) SOX2-ON re-express markers associated with pluripotency when cultured in FLC medium.

**Supplementary Figure relating to Figure 3.**
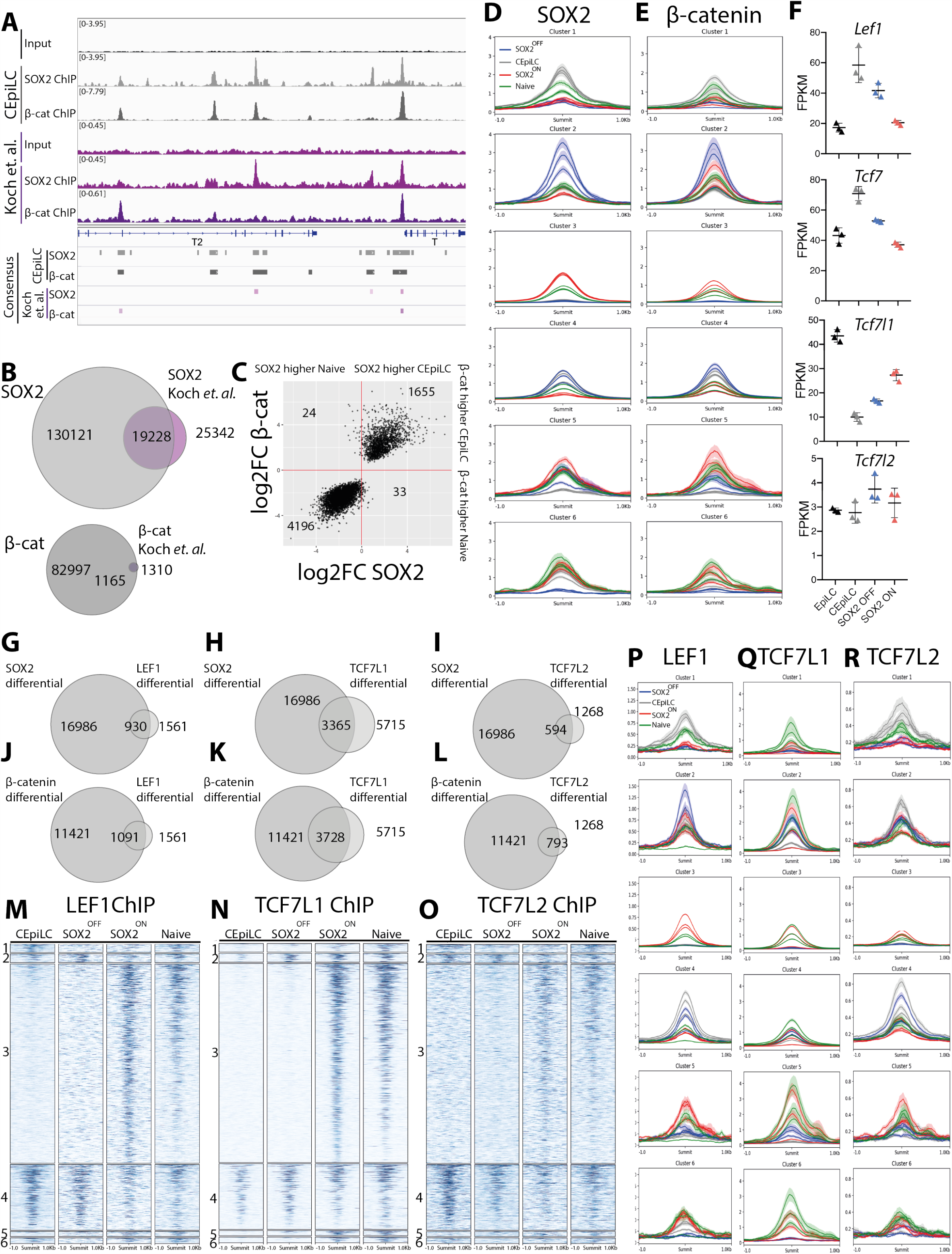
TCF/LEF are redistributed with SOX2 and β-catenin. (A) Comparison of SOX2 and β-catenin ChIP-seq peaks identified in data from this study with (Koch et al., 2017). (B) Our consensus SOX2 and β-catenin peak sets include the majority of those identifiable from the data from (Koch et al., 2017), plus a large number of additional peaks. (C) Peaks differentially occupied by β-catenin (FDR <0.05, n = 3) between ES cells and CEpiLCs exhibit a correlated change of SOX2 occupancy. (D) Metaplots of triplicate SOX2 and (E) β-catenin ChIP-seq signals at cell-type specific SOX2 bound CREs classified in Fig 3E. (F) *Lef1, Tcf7*, and *Tcf7L1* are differentially expressed across cell-states. Triplicate FPKM values for each gene are shown. The majority of (G) LEF1, (H) TCF7L1, and (I) TCF7L2 differential peaks overlap SOX2 differential peaks. The majority of (J) LEF1, (K) TCF7L1, and (L) TCF7L2 differential peaks overlap β-catenin differential peaks. (M) LEF1, (N) TCF7L1, and (O) TCF7L2 exhibit a similar cell-type specific pattern of occupancy as β-catenin at SOX2 differential peaks (compare to Fig 3E,F). Metaplots of triplicate (P) LEF1, (Q) TCF7L1 and (R) TCF7L2 ChIP-seq signals at cell-type specific SOX2 bound peaks classified in Fig 3E. All bars in metaplots = sem.

**Supplementary Figure relating to Figure 4.**
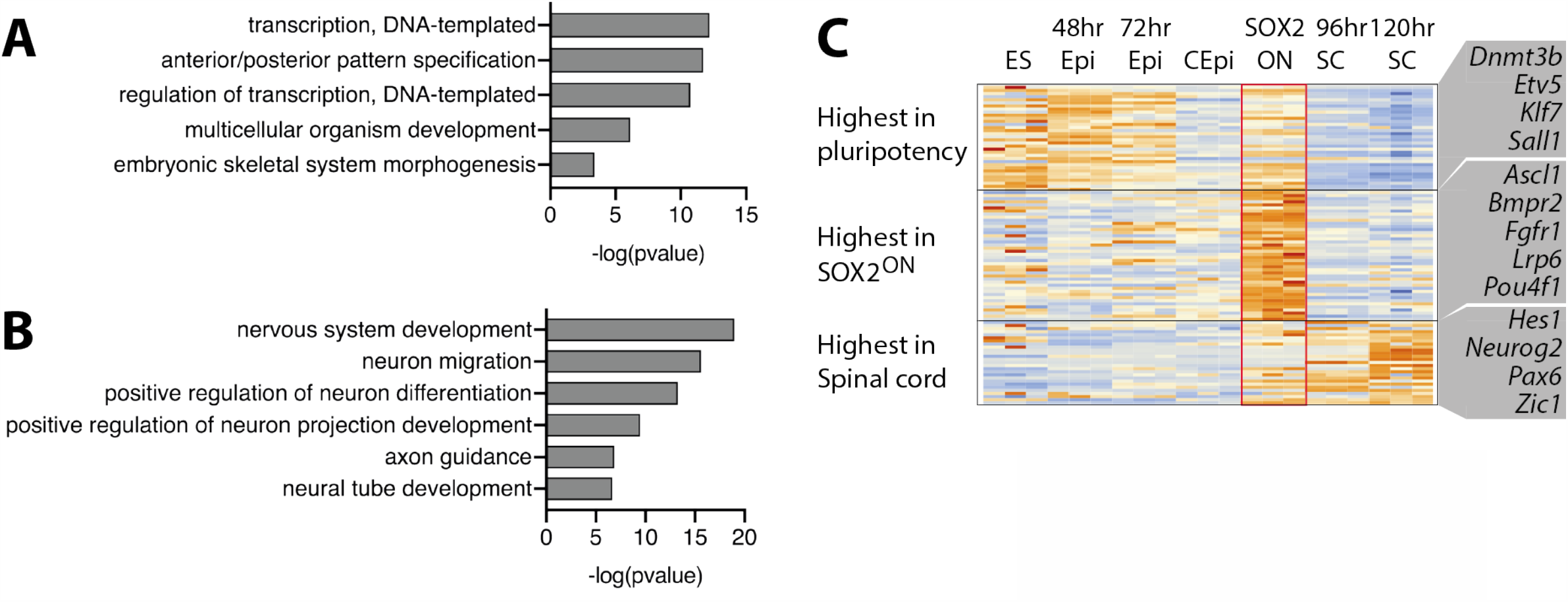
SOX2 ON express spinal-cord markers at low levels. (A) Differentially expressed genes associated with CEpiLC specific cluster 1 CREs are enriched for biological processes underlying the patterning of the anterior-posterior axis. (B) Differentially expressed genes associated with SOX2-ON specific cluster 3 CREs are enriched for biological processes related to nervous system development. (C) Cluster 3 associated genes with GO terms related to with nervous system development from (B) that are expressed at high levels in spinal-cord (SC) neural progenitors exhibit comparatively low expression in SOX2-ON, ES cells, EpiLCs and CEpiLCs. Illustrative genes for each cluster are highlighted. Spinal cord progenitor data reanalysed from (Gouti et al., 2014).

**Supplementary Figure relating to Figure 5.**
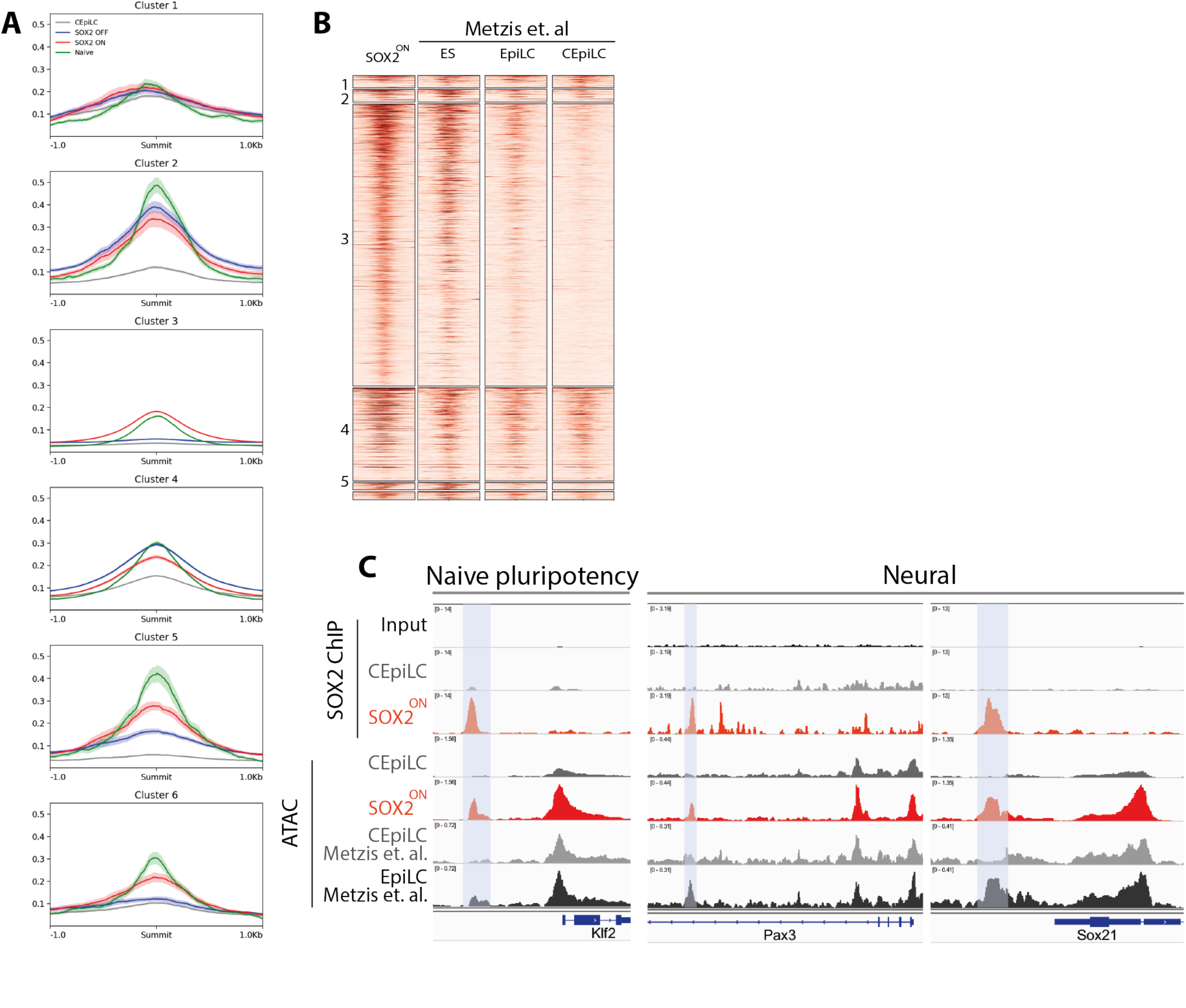
High SOX2 levels promote chromatin accessibility at pluripotency and neural genes. (A) Metaplots of ATAC-seq data at cell-type specific SOX2 bound CREs classified in Fig 3E. Naïve ATAC-seq data reanalysed from (Kearns et al., 2015). (B) ATAC-seq accessibility is greater at cluster 3 CREs in SOX2-ON, ES cells, and EpiLCs compared to CEpiLCs. Data reanalysed from (Metzis et al., 2018) where indicated. (C) Correlated SOX2 occupancy and ATAC-seq accessibility at representative cluster 3 CREs at pluripotency and neural genes from data generated in this study and (Metzis et al., 2018). Blue bars highlight peaks with greater SOX2 occupancy and ATAC-seq accessibility in SOX2-ON and EpiLCs compared to CEpiLCs.

**Supplementary Figure relating to Figure 6.**
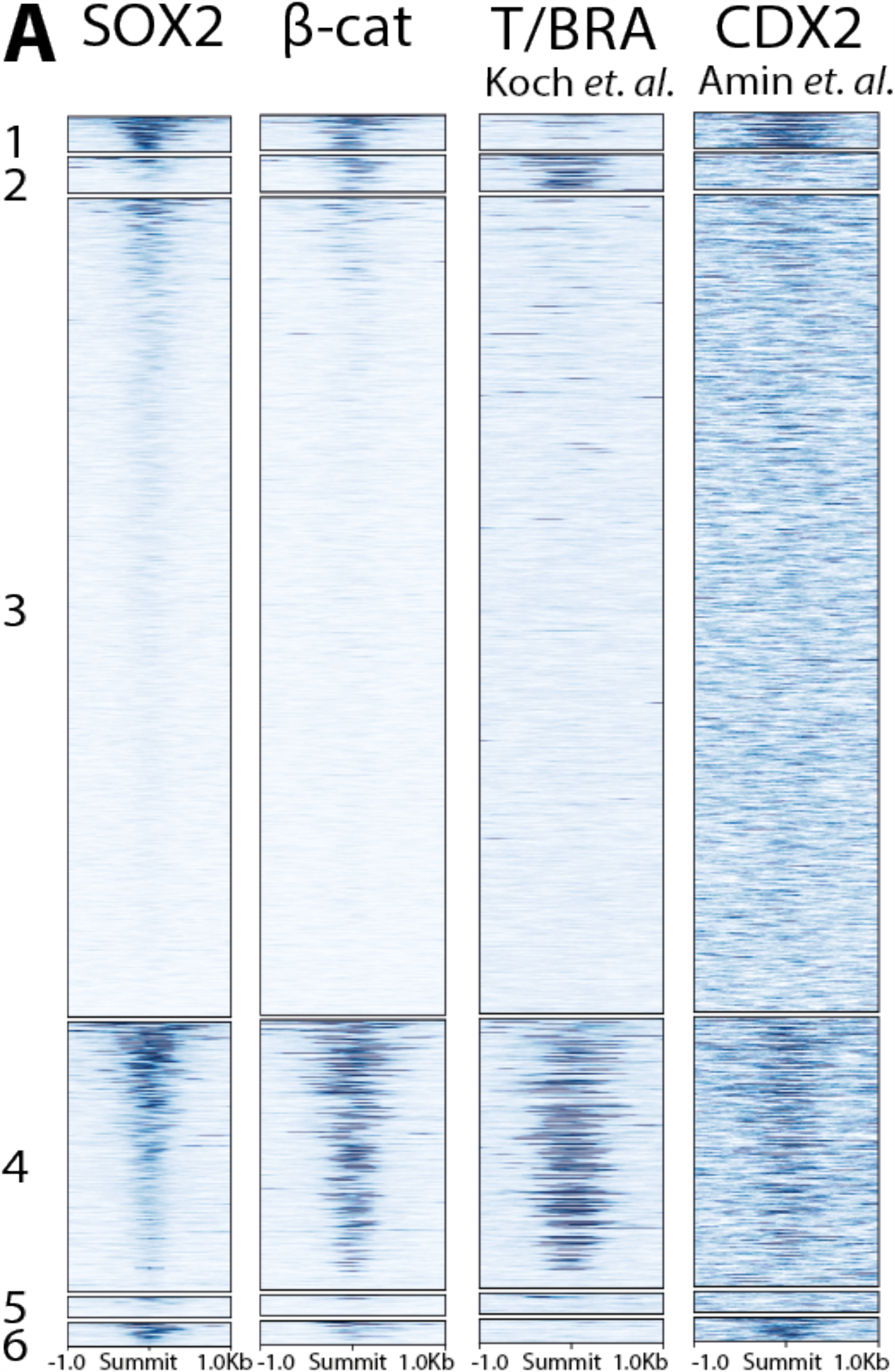
SOX2 and β-catenin co-occupy cell-type specific CREs with T/BRA and CDX2. (A) SOX2 and β-catenin occupancy is correlated with T/BRA and CDX2 in CEpiLCs at cell-type specific CREs classified in Fig 3-E. All bars in metaplots = sem.

**Supplementary Figure relating to Figure 7.**
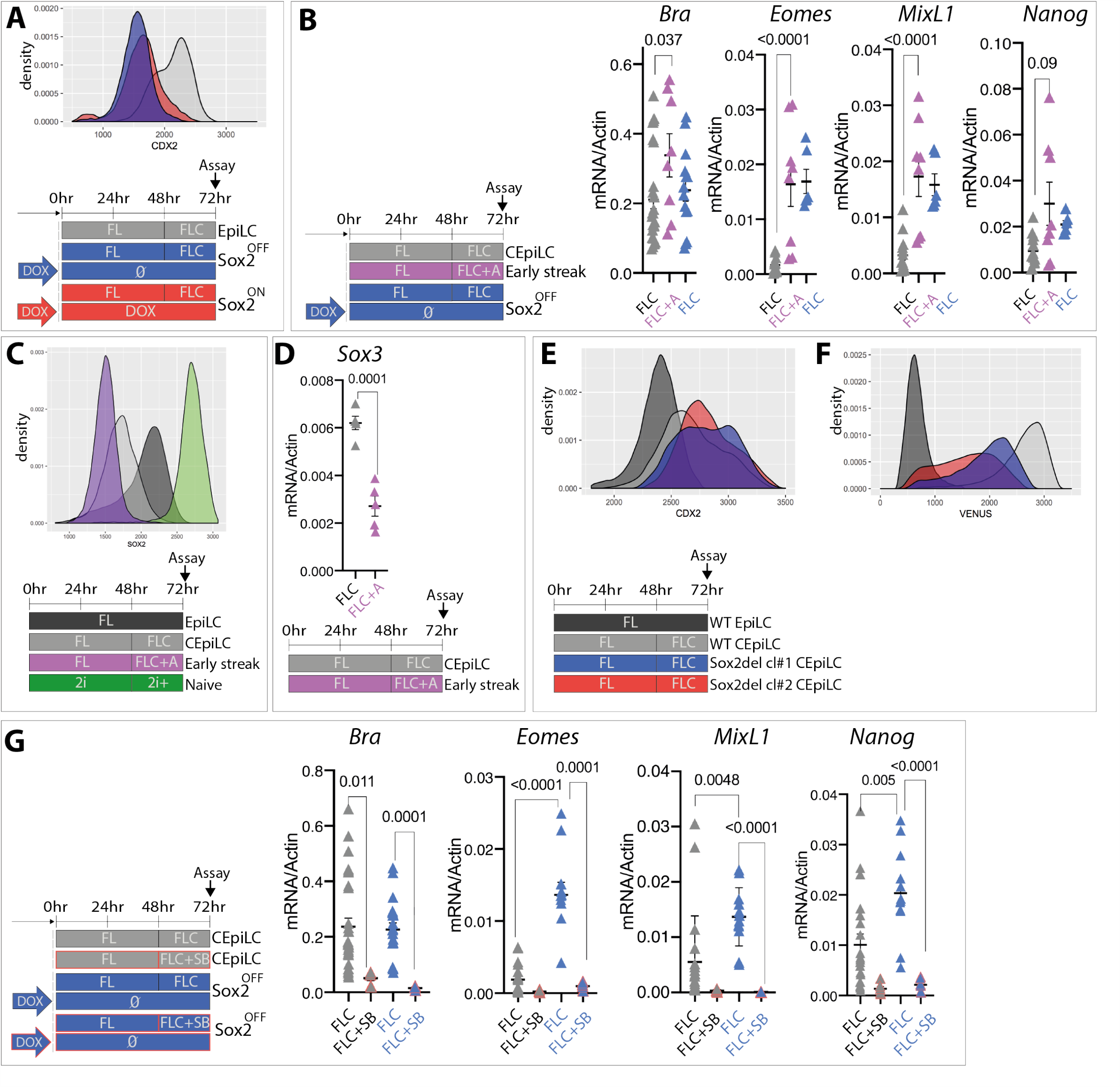
Repression of SOX2/SOX3 and Cdx2 expression in early-streak progenitors. (A) CDX2 expression is absent in SOX2-OFF and SOX2-ON cultured in FLC medium. (B) Expression of primitive streak markers in CEpiLCs and SOX2 OFF cultured in FLC medium, and early-streak progenitors induced by the addition of activin (10ng/ml) to FLC medium. (C) SOX2 levels in EpiLCs, CEpiLCs, pluripotent and early streak progenitors. CHIR concentration was increased to 5M in 2i+ conditions. (D) Sox3 levels are reduced in early-streak progenitors induced by the addition of activin to FLC medium. (E) CDX2 expression is similar, whereas (F) VENUS fluorescence is reduced in two Sox2del subclones analysed in Fig 6F compared to parental the CDX2 Intron reporter line. (G) Primitive streak marker expression is reduced in CEpiLCs and SOX2 OFF cultured in FLC medium plus SB431542. Individual biological replicates are plotted, and p-values are show.

## Materials and Methods

### Cell lines

All cell lines were maintained and experiments performed at 37°C with 5% carbon dioxide (CO_2_). All ESC lines used were derived from the XY HM1 TetON line (Serafimidis et al., 2008) which was used as the WT control. Sox2 TetON, *Cdx2* intron CRE reporter, and *Cdx2* intron CRE reporter Sox2del lines were generated as described below. Cell lines were validated by DNA sequencing and flow cytometry, and routinely tested for mycoplasma.

### Sox2 TetON

Sox2 TetON was generated by introducing a silent G>A mutation 54bp into the ORF of Sox2 cDNA using site directed mutagenesis to ablate the PAM site targeted by the guide Sox2_crispr_1 (see key reagents table). The Sox2_crispr_1 -insensitive Sox2 cDNA was then subcloned into pBI2 using Sal1/Not1 restriction digest and subsequently cloned into the HPRT locus targeting vector Hprt2 as described previously by (Serafimidis et al., 2008). The HPRT_TetON-SOX2_54G>A construct was electroporated into HM1 TetON ES cells and integrants were selected by culturing for 10 days in HAT containing ES medium. Construct integration was confirmed by genotyping, and transgene expression was confirmed by flow cytometry. SOX2_54G>A targeted cells were then adapted to 2i culture, the transgene was induced with Doxycycline (1μg/ml), and cells were electroporated with Sox2_crispr_1 using an Amaxa Nucleofector to ablate endogenous Sox2 gene expression. Electroporated cells were seeded at clonal density on gelatin plates in 2i medium and the following day were selected using puromycin (1μg/ml) for 36 hrs. Resistant clones were grown, picked, expanded, and screened for Sox2 expression by flow cytometry to detect complete loss of SOX2 expression following withdrawal of Dox. Following ablation of endogenous SOX2, Sox2 TetON were stably maintained in pluripotency conditions by addition of 1μg/ml Dox to Serum + LIF ES culture medium.

### *Cdx2* intron CRE reporter

GeneBlock oligonucleotides coding for a 1.6kb fragment of WT sequence comprising the SOX2 peaks within CDX2 intron 1, or the same region in which 7 predicted Sox2 motifs (JASPAR) (Khan et al., 2018) were scrambled, were cloned by Gibson assembly into a pENTR11 backbone upstream of an hsp68 minimal promoter driving expression of a Venus-H2B transgene. WT and scrambled sequences are detailed in ‘key reagents’ section. Additionally, the Gateway cassette from FuTetO-GW (Addgene) was cloned by Gibson assembly into the Asc1/Pme1 site of Hprt2 (Serafimidis et al., 2008) to yield HPRT_GW. LR clonase (Invitrogen) was used to induce recombination between the pENTR reporter construct and HPRT_GW, yielding HPRT-locus targeting constructs which were used to generate stable lines in HM1 TetON ES cells as described for Sox2 TetON.

### ES Cell culture and differentiation

All mouse ESCs were propagated on mitotically inactivated mouse embryonic fibroblasts (feeders) in DMEM knockout medium supplemented with 1000U/ml LIF (Chemicon), 10% cell-culture validated fetal bovine serum, penicillin/streptomycin, 2mM L-glutamine (GIBCO). To obtain EpiLCs and CEpiLCs, ESCs were differentiated as previously described (Gouti et al., 2014) with the addition of the porcupine inhibitor LGK974 (Cayman) in all culture medium. Briefly, ESCs were dissociated with 0.05% trypsin, and plated on tissue-culture treated plates for two sequential 20-minute periods in ESC medium to separate them from their feeder layer cells which adhere to the plastic. To start the differentiation, cells remaining in the supernatant were pelleted by centrifugation, counted, and resuspended in N2B27 medium containing 10ng/ml bFGF (Peprotech) + 5μM LGK974, and 50,000 cells per 35mm gelatin-coated CELLBIND dish (Corning) were plated. N2B27 medium contained a 1:1 ratio of DMEM/F12:Neurobasal medium (GIBCO) supplemented with 1xN2 (GIBCO), 1xB27 (GIBCO), 2mM L-glutamine (GIBCO), 40mg/ml BSA (Sigma), penicillin/streptomycin and 0.1mM 2-mercaptoethanol. To generate EpiLCs, the cells were grown for 72 hrs in N2B27 + 10ng/ml bFGF + 5μM LGK974 (FL-medium). To generate CEpiLCs, cells were cultured with N2B27 + 10ng/ml bFGF + 5μM LGK974 for 48 hrs, then N2B27 + 10ng/ml bFGF + 5uM LGK974 + 5mM CHIR99021 (Axon) (FLC-medium) for a further 24hrs (day 3 in Gouti et al., 2014). CEpiLCs were differentiated to spinal cord neural progenitors by removal of bFGF and CHIR from culture medium at 72hrs, and to paraxial mesoderm by removal of bFGF and maintenance of 5mM CHIR from 72hrs onwards. When investigating the activity of Nodal signalling, either 10ng/ml recombinant activin (R&D) or 10uM ALK-inhibitor SB-431542 (LKT) was included in bFGF/CHIR containing medium. Experiments conducted in 2i medium were initiated by separating serum/LIF-grown ES cells from feeders as described above and plating onto gelatin coated CELLBIND dishes in N2/B27 containing basal medium supplemented with 3μM CHIR and 500nM PD0325901. For all experiments described cells were cultured for 48hrs before changing medium. Medium changes were then made every 24 hours.

### Immunofluorescence

Cells were washed in PBS and fixed in 4% paraformaldehyde in PBS for 15min at 4°C, followed by two washes in PBS and one wash in PBST (0.1% Triton X-100 diluted in PBS). Primary antibodies were applied overnight at 4°C diluted in filter-sterilized blocking solution (1% BSA diluted in PBST). Cells were washed 3x in PBST and incubated with secondary antibodies at room temperature, for 1hr. Cells were washed 3x in PBST, incubated with DAPI for 5 min in PBS and washed twice before mounting with Prolong Gold (Invitrogen). Cells were imaged on a Zeiss Imager.Z2 microscope using the ApoTome.2 structured illumination platform. Z stacks were acquired and represented as maximum intensity projections using ImageJ software. The same settings were applied to all images. Immunofluorescence was performed on a minimum of 2 biological replicates, from independent experiments. All primary antibodies used in this study are listed in ‘key reagents’. Secondary antibodies raised in donkey coupled to AlexaFluor 488, 568, or 647 fluorophores (Molecular Probes) were used at 1:1000 dilution throughout unless otherwise stated.

### Intracellular Flow cytometry

Cells were washed in PBS and dissociated with minimal accutase (GIBCO). Once detached cells were collected into 1.5mL Eppendorf tubes by dissociating in N2B27 and pelleted. Cells were resuspended in PBS, pelleted and resuspended in 4% paraformaldehyde in PBS. Following 15min incubation at 4 C, cells were centrifuged, resuspended in PBS, and stored at 4°C for future analysis. On the day of flow cytometry, cells were counted and equal cell numbers were transferred for staining in v-bottom 96 well plates. Samples were pelleted and resuspended in 50uL FACS block (PBS + 0.2% Triton + 3% BSA). After 10min incubation at room temperature antibodies were added to the sample and incubated overnight at 4°C. Cells were pelleted at 700rcf for 5min and resuspended in 50uL FACS block. One additional wash was performed before acquisition on a Fortessa flow cytometer (BD). Analysis was performed using the R-package flowCore (Hahne et al., 2009) and data was graphed using ggplot2 (Wickham, 2016).

### RNA extraction

RNA used for qPCR or RNA-seq was extracted from cells using a QIAGEN RNeasy kit in RLT buffer, following the manufacturer’s instructions. Extracts were digested with DNase I to eliminate genomic DNA.

### cDNA synthesis, and qPCR analysis

First strand cDNA synthesis was performed using Superscript III (Invitrogen) using random hexamers and was amplified using PowerUp SYBR-Green Mastermix (Applied Biosystems). qPCR was performed using the Applied Biosystems QuantStudio Real Time PCR system. PCR primers were designed using online GenScript qPCR primer design tool. Expression values for each gene were normalized against b-actin, using the delta-delta CT method. qPCR analysis was performed on pooled samples from a minimum of 3 independent experiments for every primer pair analysed. Primer sequences are available in key reagents. Data was graphed using Prism software.

### RNA-seq

Libraries were prepared using the KAPA mRNA HyperPrep kit (Roche) and sequenced as 76bp single-end, stand-specific reads on the Illumina HiSeq 4000 platform (Francis Crick Institute).

### RNA-seq analysis

Adapter trimming was performed with cutadapt (version 1.16) (Martin, 2011) with parameters “--minimum-length=25 --quality-cutoff=20 -a AGATCGGAAGAGC”, and for paired-end data “-A AGATCGGAAGAGC” was appended to the command. The RSEM package (version 1.3.0) (Li and Dewey, 2011) in conjunction with the STAR alignment algorithm (version 2.5.2a) (Dobin et al., 2013) was used for the mapping and subsequent gene-level counting of the sequenced reads with respect to mm10 RefSeq genes downloaded from the UCSC Table Browser (Karolchik et al., 2004) on 11th December 2017. The parameters passed to the “rsem-calculate-expression” command were “--star --star-gzipped-read-file --star-output-genome-bam --forward-prob 0”, and for paired-end data “--paired-end” was appended to the command. Differential expression analysis was performed with the DESeq2 package (version 1.16.1) (Love et al., 2014) within the R programming environment (version 3.4.1). An adjusted p-value of <= 0.05 was used as the significance threshold for the identification of differentially expressed genes.

### RNAseq clustering

The R ‘kmeans’ function was used to cluster standardised (z-transformed) FPKM values across biological conditions prior to plotting with R ‘heatmap2’ function. The lowest value of k able to partition gross trends in the data was chosen.

### RNAseq-associating differential gene expression with differential SOX2 occupancy

Homer ‘annotatePeaks.pl’ was used to associate consensus SOX2 ChIP peaks with nearest gene promoters. SOX2 associated genes were then filtered based upon their differential expression in pairwise comparisons between CEpiLC, SOX2 OFF, and SOX2 ON using Deseq2. Mean FPKM values from triplicate samples were z-transformed across the three experimental conditions to standardise fold-change in expression and plotted using ggplot2.

### RNAseq GO enrichment

The online functional annotation tool of the DAVID bioinformatics resource https://david.ncifcrf.gov/summary.jsp was used with default parameters to identify statistically enriched biological process annotations within sets of transcript and gene IDs, and to calculate associated Benjamini-hochberg adjusted p-values.

### RNAseq-comparison of in vitro to in vivo epiblast differentiation

Principal component analysis was performed on mRNAseq data from duplicate 2i, and triplicate ICM, E4.5 epiblast and E5.5 epiblast samples from Boroviak et al using using the R function prcomp. PC1 aligned with developmental time, whereas PC2 separated in vitro (2i) and in vivo (ICM, E4.5 and E5.5) derived samples. The top 300 genes contributing most positively and negatively to PC1 were selected to represent the gene expression dynamics observed to occur during epiblast differentiation in vivo, and the dynamics of their expression during in vitro differentiation of ESCs to EpiLCs was represented by plotting standardised (z-transformed) FPKM values using heatmap2.

### ChIP-seq

Adherent cells were washed 3 times with PBS, fixed with gentle agitation for 45min at room temperature with fresh 2mM Di(N-succinimidyl) glutarate (Sigma) in PBS, washed an additional 3 times with PBS, then fixed for 10min at room temperature with 1% molecular biology grade PFA in PBS. Fixation was quenched by addition of 250mM glycerine for 5 minutes, followed by additional washing with PBS. Plates were cooled, cells were scraped into tubes in a low volume of PBS 0.02% Triton X-100, and pelleted by centrifugation at 1000rpm for 5min at 4oC before snap freezing in liquid nitrogen and storing at −80°C. Approximately 5*10^6 cells were transferred to a Diagenode TPX tube and resuspended in ice cold shearing buffer (see key reagents) containing 0.3% SDS and protease inhibitors (Sigma). Chromatin was sheared using a Bioruptor plus: 25 cycles of 30s On/30s Off on high setting, and lysates were then diluted to 0.15% SDS and cleared by centrifugation at 14,000rpm for 10min at 4°C. 1/20^th^ of the chromatin from ∼1*10^7 cells was set aside and frozen for subsequent use as input control, and the remainder was incubated overnight at 4°C under rotation with 100ul of protein G dynabeads (Invitrogen) pre-loaded for 4hrs at RT with 5ug of ChIP antibodies diluted in shearing buffer containing 0.15% SDS (see key reagents). Beads were magnetically immobilised, unbound supernatant discarded, and beads were sequentially washed under rotation twice with Wash Buffer 1, once with Wash Buffer 2, once with Wash Buffer 3, and twice with Wash Buffer 4 for 5 minutes each, magnetically capturing beads between each wash (see key reagents for composition of buffers). Chromatin was eluted from beads by incubating twice at 65°C for 10mins in 100ul elution buffer on a shaking heat block, capturing beads between each elution step and then pooling each eluted fraction. Input samples were made up to 200ul with elution buffer, 6.4ul of 5M NaCl was added to each input or IP sample, and all samples were de-crosslinked overnight at 65°C. Samples were incubated for 2hrs at 37°C with 0.2ug/ml PureLink RNAse A (Invitrogen), then supplemented with 5mM EDTA and incubated for an additional 2hrs at 45°C with 0.2ug/ml proteinase K (Thermo Scientific) before purifying DNA with Qiagen PCR cleanup columns. DNA fragmentation of IP and input samples was confirmed by Agilent TapeStation prior to library preparation using NEB Ultra II DNA. Biological triplicates were obtained for all conditions from separate experiments. Libraries were sequenced as single-end, 76bp reads on the Illumina High-Seq 4000 platform (Francis Crick Institute).

### ChIP-seq analysis

The nf-core/ChIP-seq pipeline (version 1.1.0; https://doi.org/10.5281/zenodo.3529400) (Ewels et al., 2020) written in the Nextflow domain specific language (version 19.10.0) (Di Tommaso et al., 2017) was used to perform the primary analysis of the samples in conjunction with Singularity (version 2.6.0) (Kurtzer et al., 2017). The command used was “nextflow run nf-core/ChIP-seq --input design.csv --genome mm10 --gtf refseq_genes.gtf --single_end --narrow_peak --min_reps_consensus 2 -profile crick -r 1.1.0”. To summarise, the pipeline performs adapter trimming (Trim Galore! - https://www.bioinformatics.babraham.ac.uk/projects/trim_galore/), read alignment (BWA) (Li and Durbin, 2009) and filtering (SAMtools) (Li et al., 2009); (BEDTools) (Quinlan and Hall, 2010); (BamTools) (Barnett et al., 2011); (pysam - https://github.com/pysam-developers/pysam); (picard-tools - http://broadinstitute.github.io/picard), normalised coverage track generation (BEDTools) (Quinlan and Hall, 2010); (bedGraphToBigWig) (Kent et al., 2010), peak calling (MACS) (Zhang et al., 2008) and annotation relative to gene features HOMER) (Heinz et al., 2010), consensus peak set creation (BEDTools) (Quinlan and Hall, 2010), differential binding analysis (featureCounts) (Liao et al., 2014); (DESeq2) (Love et al., 2014) and extensive QC and version reporting (MultiQC) (Ewels et al., 2016); (FastQC - https://www.bioinformatics.babraham.ac.uk/projects/fastqc/); (preseq); deepTools (Ramírez et al., 2016); (phantompeakqualtools) (Landt et al., 2012). All data was processed relative to the mouse UCSC mm10 genome (UCSC) (Karolchik et al., 2004) downloaded from AWS iGenomes (https://github.com/ewels/AWS-iGenomes). Peak annotation was performed relative to the same GTF gene annotation file used for the RNA-seq analysis. Tracks illustrating representative peaks were visualised using the IGV genome browser (Robinson et al., 2011).

### ChIP-seq peak clustering

SOX2 peaks were manually assigned to 6 clusters based upon differential occupancy between WT, ‘SOX2 OFF’, and ‘SOX2 ON’ samples. Peaks in clusters 1, 2 and 3 had the highest mean read counts across biological triplicate samples in either WT, ‘SOX2 OFF’, or ‘SOX2 ON’ respectively, and were statistically different (FDR <0.05) as determined by DeSeq2 in comparison to all other experimental conditions. Cluster 4, 5, and 6 peaks were statistically different to only one of the other experimental conditions. Genomic intervals defined by SOX2 peaks within these clusters were used to plot metaplots and heatmaps from the BigWig files generated from the nf-core/ChIP-seq and nf-core/ATAC-seq pipelines using deepTools.

### ChIP-seq Motif enrichment

Motifs enriched within each SOX2 peak cluster were identified using Homer (Heinz et al., 2010) findMotifsGenome using default parameters. Region size = 200b (+/-100bp adjacent to peak centre).

### ChIP-seq Motif scoring (FIMO)

Regions of +/-100bp adjacent to SOX2 ChIP-seq peak centres were used as inputs for the motif scanning tool http://meme-suite.org/tools/fimo (Bailey et al., 2009). The Sox2 motif MA0143.3 (JASPAR) (Khan et al., 2018) was used as a target. p-value threshold was set to p <0.1 so as to include low scoring SOX2 motifs present within peak sets.

### ChIP-seq peak intersection

BEDtools (Quinlan and Hall, 2010) intersectBed was used to identify genomic intervals overlapping by > 10% in.bed files listing coordinates of consensus and differentially occupied peak sets for each immunoprecipitated factor.

### ATAC-seq

ATAC-seq sample preparation was performed as described in Metzis et al. Libraries were sequenced as paired-end, 101bp reads on the Illumina High-Seq 4000 platform (Francis Crick Institute).

### ATAC-seq analysis

The nf-core/atacseq pipeline (version 1.0.0; https://doi.org/10.5281/zenodo.2634133) (Ewels et al., 2020) written in the Nextflow domain specific language (version 19.10.0) (Di Tommaso et al., 2017) was used to perform the primary analysis of the samples in conjunction with Singularity (version 2.6.0) (Kurtzer et al., 2017). The command used was “nextflow run nf-core/ATAC-seq --design design.csv --genome mm10 --gtf refseq_genes.gtf -profile crick -r 1.0.0”. The nf-core/ATAC-seq pipeline uses similar processing steps as described for the nf-core/ChIP-seq pipeline in the previous section but with additional steps specific to ATAC-seq analysis including removal of mitochondrial reads.

### Nucleosome analysis

The NucleoATAC package (version 0.3.4) (Schep et al., 2015) was run in default mode. Analysis was performed on all genomic intervals called as peaks from ATAC-seq data as described above. Metaplots of the occ.bedgraph files for each experimental condition were plotted using deepTools to score the average nucleosome occupancy within each peak cluster. Tracks of the occ.bedgraph and nucleoatac_signal.smooth.bedgraph files were visualised using the IGV genome browser (Robinson et al., 2011) to illustrate the occupancy and position of nucleosomes at genomic intervals of interest.

### Statistical analysis

Statistical analysis and software are described in figure legends and key reagents and software

### Data availability

The GEO accession number for the next-generation sequencing data reported in this paper is GSE162774. Reanalysed data samples are listed in key reagents and software.

